# Greenscreen decreases Type I Errors and increases true peak detection in genomic datasets including ChIP-seq

**DOI:** 10.1101/2022.02.27.482177

**Authors:** Sammy Klasfeld, Doris Wagner

## Abstract

Chromatin immunoprecipitation followed by sequencing (ChIP-seq) is used widely to identify both factor binding to genomic DNA and chromatin modifications. Analysis of ChIP-seq data is impacted by regions of the genome which generate ultra-high artifactual signals. To remove these signals from ChIP-seq data, ENCODE developed blacklists, comprehensive sets of regions defined by low mappability and ultra-high signals for human, mouse, worm, and flies. Currently, blacklists are not available for many model and non-model species. Here we describe an alternative approach for removing false-positive peaks we called “greenscreen”. Greenscreen is facile to implement, requires few input samples, and uses analysis tools frequently employed for ChIP-seq. We show that greenscreen removes artifact signal as effectively as blacklists in Arabidopsis and human ChIP-seq datasets while covering less of the genome, dramatically improving ChIP-seq data quality. Greenscreen filtering reveals true factor binding overlap and of occupancy changes in different genetic backgrounds or tissues. Because it is effective with as few as three inputs, greenscreen is readily adaptable for use in any species or genome build. Although developed for ChIP-seq, greenscreen also identifies artifact signals from other genomic datasets including CUT&RUN. Finally, we present an improved ChIP-seq pipeline which incorporates greenscreen, that detects more true peaks than published methods.

**One Sentence Summary:** A facile method for removing artifact signal from ChIP-seq that improves downstream analyses

## Introduction

Chromatin immunoprecipitation followed by sequencing (ChIP-seq) probes the association of a factor or modification with chromatin^1^. After factor crosslinking to chromatin and shearing of the genomic DNA, DNA fragments associated with the factor of interest are enriched by immunoprecipitation and sequenced after crosslink reversal^1^. It has been established that ChIP-seq produces both true-positive signals as well as artifactual signal in the process of sequence enrichment ^2–6^.

Guidelines for accurate analyses of ChIP-seq suggest using experimental controls, “input” DNA or “mock” ChIP, to account for areas of the genome with sequencing efficiency biases^2–7^. “Input” control samples are identical to a given experimental sample except that they were not subjected to immunoprecipitation^1^. “Mock’’ samples are ChIP-seq reactions where either the genetic background lacks the antigen or antiserum is employed that does bind the antigen ^2–7^. In the absence of sequencing biases, “input” DNA should appear uniformly distributed across the genome, while no peaks are expected for the “mock” ChIP experiment ^2,3^.

However, some regions in the genome give rise to amplified artifact signals that are not efficiently removed through normalization of experimental controls and impact experimental analysis ^3,8–10^. These ultrahigh signals are present in ChIP, mock, and input samples at various levels, and failure to remove the artifact signal prevents accurate analysis of sample quality, replicate concordance, and identification of factor binding sites ^8–10^. To identify and mask out these regions from downstream analysis, ENCODE curated a filter called “blacklist” for mouse, human, fly and worm ^7–10^.

Often artifacts occur near assembly gaps and in other genomic regions with low copy repeat elements and have high ratios of multi-mapped to unique reads ^9–10^. The genomic regions that give rise to artifact signals are invariant for a given species with regards to developmental stage/tissue, yet the signal strength in these regions can vary between experiments ^8–11^. Where these signals arise is also sensitive to the genome build, as new artifact prone genomic regions are added and others are lost due to added/fixed gaps, inclusion of centromeric regions or additional satellite sequences. Therefore, new blacklists have been generated for successive genome assemblies ^10^.

The blacklist filter identifies areas of the genome which have low mappability rates or contain high artifact signals. A software called UMap^12^ uses genome assembly files to measure a region’s mappability, a metric for how uniquely all predicted read-length fragments map to the genome. In addition, Blacklists identify artifact signal in inputs (top 0.1% given quantile-normalization of read depth) ^10^. Next, these high signal regions are merged within a certain distance (20 kb for human and 5 kb for Drosophila blacklists), if the merged region maintains an overall average signal intensity in the top 1% ^10^. ENCODE blacklists were generated using several hundreds of inputs and it was recommended that users employ these curated blacklist regions to mask out reads that overlap with them before applying ChIP-seq peak-calling software such as MACS2 ^10,13^.

However, blacklists are not available for most species. In addition, the blacklist generation pipeline requires tools that are not frequently used in ChIP-seq analysis and require considerable amounts of RAM and disc storage (minimum requirements are RAM: 64+ GB; CPU: 24+ cores, 3.4+ GHz/core; https://github.com/Boyle-Lab/Blacklist) ^10^. Finally, existing blacklists for *H. sapiens*, *M. musculus*, *D. melanogaster* and *C. elegans* employed hundreds of inputs ^9–10^. Given that such large input numbers are not available for most other model or non-model species, there is a need for a facile tool that allows identification of artifact signal regions with few inputs.

To address this need, we developed an alternative approach for removing ultra-high signal peaks from ChIP-seq datasets that we called “greenscreen”. We hypothesized that we could identify regions of ultra-high noise from a small number of inputs with a common peak-calling tool, MACS2 (2.2.7.1)^13^. We show here that our method is robust with as few as three inputs and performs as well as blacklisting in masking artificial signals from Arabidopsis and human ChIP-seq datasets. Because of these attributes, and because it utilizes software that is frequently used for ChIP-seq peak calling, greenscreen can readily be applied to any model and non-model species. In addition to increased versatility and ease of implementation, greenscreen masks less of the genome and fewer genes than blacklists. Greenscreen uncovers true ChIP-seq replicate concordance, true factor binding overlaps or occupancy changes in different conditions. We developed a new ChIP-seq pipeline that incorporates greenscreen and identifies a larger number of true peaks than published methods.

## Results

### Development of the greenscreen mask

To design artificial signal masks, we first focused on *Arabidopsis thaliana,* a plant model organism for which there are currently no blacklists available. We collected ChIP inputs and identified twenty that passed our quality control (see methods). These inputs were derived from different tissues and generated in different laboratories (Table S1). Ultrahigh signal peaks were present in these inputs (Figure S1A) as previously observed for mammals, flies and worms ^8–10^.

Next, we generated a blacklist for *Arabidopsis thaliana* using UMap (1.1.0) ^12^ on the TAIR10 *Arabidopsis thaliana* genome assembly ^14^ as a positive control. Employing the twenty inputs in Table S1 and the UMAP output, we applied the recently published blacklist tool (Figure S2) ^10^. We manually adjusted the merge parameter for regions with artifact signal to 5 kb as for the Drosophila blacklist, since Arabidopsis and Drosophila have similar genome sizes. As previously described, we removed reads that overlapped with blacklist regions prior to calling ChIP-seq peaks (Figure 1A, Figure S2). The resulting Arabidopsis thaliana blacklist masked 2.82% of the genome, which includes 268 protein coding genes (Table 1).

**Figure 1.**
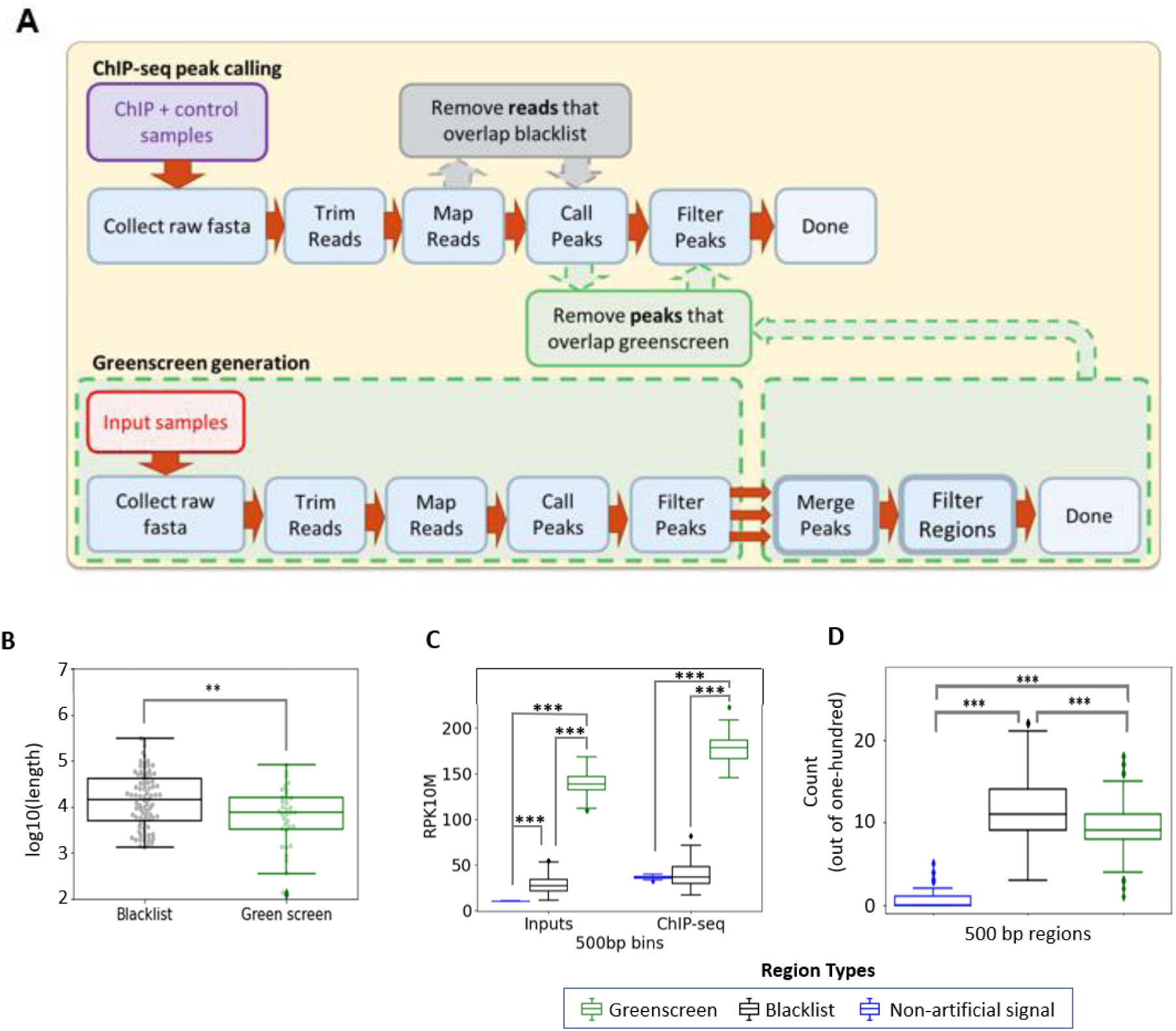
Generating ultrahigh signal blacklist and greenscreen masks for Arabidopsis. (A) ChIP-seq analysis workflow (top). Generating a greenscreen mask from input (bottom). Peaks called in all inputs are concatenated (three-red arrows) and those within a set distance are merged into a shared region. Regions which contain peaks in less than half the given inputs are removed. The resulting greenscreen mask is used to remove ChIP-seq peaks that overlap with it. (B) Box-and-whisker/swarmplot of artificial signal regions lengths [log10(base pairs)], in the blacklist masks (n=83, mean=4.1) and the greenscreen mask (n=36, mean=3.8). Two sample one-sided t-test **p = 1.6 e-3 (C) Grouped box-and-whisker plot of average normalized read signals (RPKM) in ChIP-seq ^17,19–24^ and input peaks that do not overlap with greenscreen of blacklist regions (blue) overlap with blacklist (black) or overlap with greenscreen (green). Bootstrapping (n=100) of five-hundred random non-overlapping regions (500bp in length). Non-artifact (blue; Input mean: 10.3; ChIP-seq mean: 36.2), blacklist (black; Input mean: 28.0; ChIP-seq mean: 38.7), or greenscreen (green; Input mean: 138.8; ChIP-seq mean: 178.3). Kruskal-Wallis H-test for difference among the three input groups (p=1.9e-58) or ChIP groups (p=5.1e-44). Gray bars above the boxplots show one-sided Mann-Whitney U rank test comparisons with Holms multiple test correction. Inputs: non-artificial signal regions and blacklist (***p=3.8e-34) or greenscreen (***p=3.8e-34), greenscreen relative to blacklist (***p=3.8e-34). ChIP-seq samples: non-artificial signal regions and blacklist (p=0.37) or greenscreen (***p=3.8e-34) and greenscreen relative to blacklist (*** p=3.8e-34). (D) Box-and-whisker plot of frequency in which one hundred randomly sampled 500bp regions from non-artifact, blacklist or greenscreen sites respectively are near (<1kb) assembly gaps. Non-artifact (mean=0.5), blacklist regions (mean=11.5), and greenscreen regions (mean=9.4). n=1000 trials with replacement were conducted. ANOVA test for difference among the three groups p = 0. Welch’s one-sided t-test with Holm’s correction relative to non-artifact regions: blacklist ***p = 0, greenscreen, ***p = 0. Welch’s two-sided t-test blacklist compared to greenscreen ***p = 1.8 e-52.

**Table 1.** Comparison of blacklist and greenscreen filter regions. All filters for artifactual signal removal were generated in this study, expect for the human blacklist which is published ^10^.

|  | **Arabidopsis**<br>**greenscreen** | **Arabidopsis**<br>**blacklist** | **Human**<br>**greenscreen** | **Human**<br>**blacklist** |
| --- | --- | --- | --- | --- |
| **Percent of**<br>**genome masked** | 0.41 | 2.49 | 0.01 | 7.36 |
| **Protein coding**<br>**genes masked** | 26 | 172 | 20 | 3390 |

To design the *Arabidopsis* greenscreen mask, we employed MACS2 ^13^, a tool routinely used to identify ChIP-seq peaks. Following the same steps used to identify peaks in ChIP-seq experiments, we trimmed, mapped, and applied MACS2 to each of the twenty inputs using the entire genome as background (Figure 1A). The broad peak setting was applied to call peaks in each of the input samples (Figure 1A). We optimized the threshold of significance for the mean q-value for each base pair in input peaks such that the resulting greenscreen filter maximizes overlap between a ChIP-seq and an orthogonal ChIP-chip dataset for the same transcription factor unlikely to be subject to these artifacts. At the same time, we strove to minimize the genomic regions and genes masked out (Figure 1A, Table 1). Using these criteria, MACS2 q<10^-10^ was chosen as the cutoff to identify input peaks. MACS2-called regions in each input were concatenated into a single list and, to maximize artifact removal, regions within a certain distance were merged (three red arrows in Figure 1A). Using the same criteria as described above for q-value cutoff, we selected the merge distance. Merging regions within 5 kb was optimal; it gave a slightly higher overlap with the orthogonal ChIP-chip dataset then the 2.5 kb merge, which also performed well (Table S2). Finally, to minimize over masking, we restricted the greenscreen mask to regions where significant input peaks were called in at least half (i.e ten) of the samples (see methods for details; Figure 1A). Our final greenscreen masks ∼0.48% of the genome and covers 86 protein coding genes (Table 1). Most of the greenscreen regions (99.9%) overlap with the blacklist (Figure S3). Because greenscreen covers less of the genome than blacklist, we mask peaks rather than reads to prevent MACS2 from calling ChIP peaks on the outer edges of artifact regions.

As expected, both the blacklist and greenscreen filter regions overlap with ultra-high signal in inputs and ChIP datasets (Figure S1A). Consistent with the higher genome coverage of the blacklist filter, blacklist regions were found to be significantly broader than greenscreen regions (Figure 1B). We next measured the read signal strength across blacklist and greenscreen regions using a bootstrap method. Five-hundred random 500 bp blacklist and greenscreen regions were chosen and measured iteratively one-hundred times in ChIP-seq and Input samples. On average the inferred read signal was significantly lower in blacklist regions than in greenscreen regions (Figure 1C). Thus, the blacklist may over mask leading to potential false negatives (Figures S1B-D).

Like blacklist regions developed for human samples, Arabidopsis artifact signals are frequently found near assembly gaps ^8–10^ (Figure S1). To determine how frequently blacklist or greenscreen regions are found within 1 kb of an assembly gap, we applied bootstrapping to one hundred random 500 bp regions from greenscreen, blacklist, and non-artificial signal regions queried iteratively a thousand times. Both blacklist and greenscreen regions were significantly more likely to be located near assembly gaps than random, non-artifact generating genomic regions (Figure 1D).

### Efficacy of artifact signal removal by greenscreen

Prior studies identified metrics suitable to assess artifact signal removal from ChIP-seq and control samples. One is the Standardized Standard Deviation (SSD), which measures the variation in signal across the genome ^9,15^. Since SSD is calculated before peak calling, we calculated each input’s SSD with ChIPQC (1.26.0) ^16^ before and after removing reads that overlap greenscreen or blacklist regions. Both greenscreen and blacklist significantly reduced the input SSD scores to the expected value of ∼1 (Figure 2A). Thus, despite covering less of the genome, the greenscreen mask is as effective as the blacklist at removing regions of strong artificial signal. We also compared the efficacy of the Arabidopsis blacklist and greenscreen in removing artifact signals from ChIP-seq replicates by testing for a reduction in the SSD ^9,15^. Applying the greenscreen filter to ChIP-seq replicates caused the SSD values to decrease with similar efficiency as filtering out reads with the Arabidopsis blacklist (Figure 2B) ^17^.

**Figure 2.**
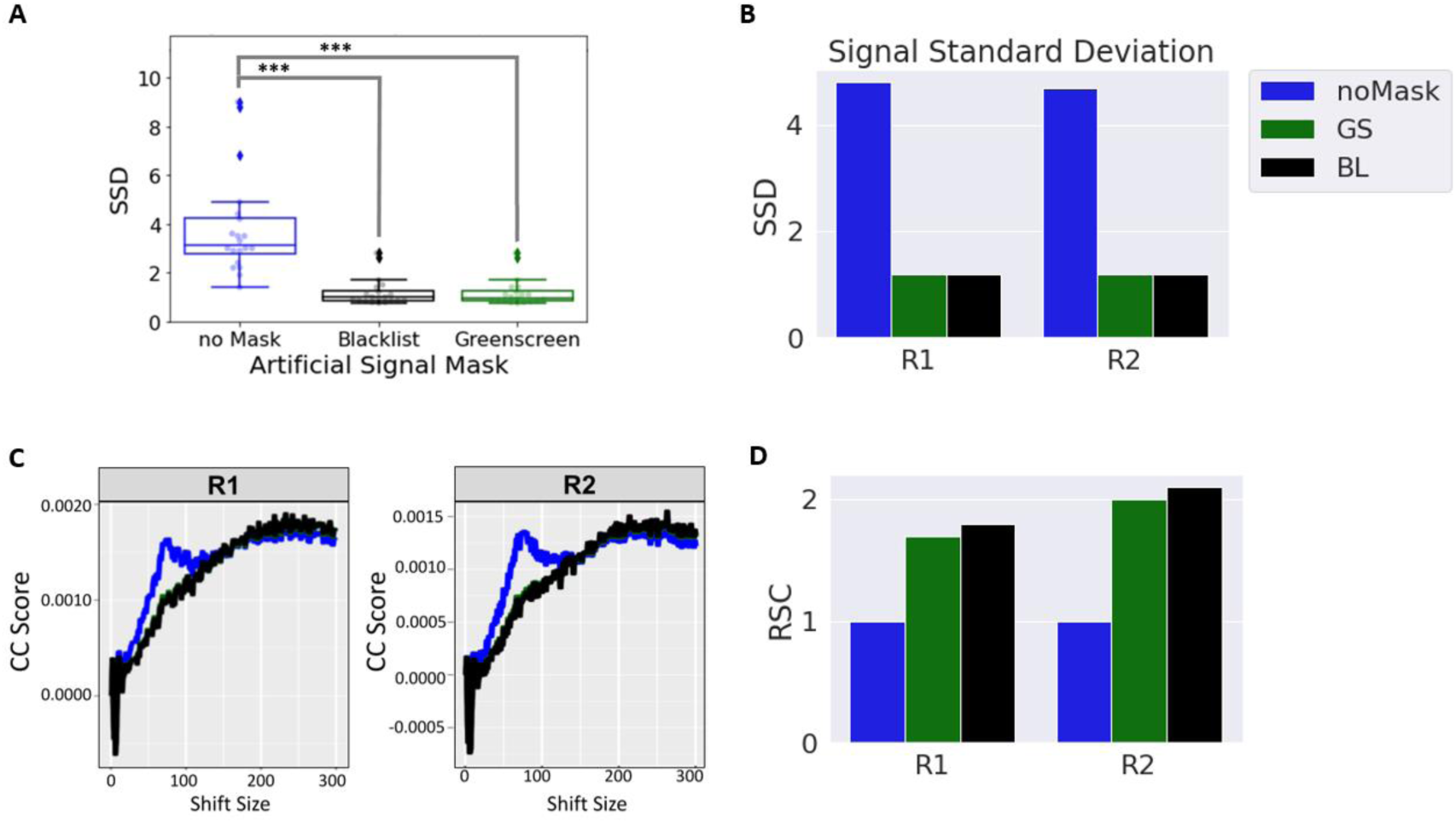
Efficacy of greenscreen artifact removal. (A) Box-and-whisker plot/swarmplot of Standardized Standard Deviation (SSD) in twenty inputs. All reads (mean=3.9), after applying blacklist (mean=1.2), and after applying greenscreen (mean=1.2). Kruskal-Wallis H-test for difference among the three groups: ***p =2.9 e-8. p-values (α=0.001). One-sided Mann Whitney U rank test with Holms correction relative to samples without mask: blacklist ***p = 5.8 e-07, greenscreen ***p = 5.8 e-07 and two-sided Mann Whitney U rank test with Holms multiple test correction blacklist relative to greenscreen: NS (not significant, p=0.9) (B) Bar graphs of Standardized Standard Deviation (SSD) values of two ChIP-seq replicates^17^ after blacklist (black) or greenscreen (green) read mask application relative to the no mask control (blue). (C) SCC curves of two ChIP-seq replicates (GSE141894)^17^. Without an artificial signal mask (blue), a phantom peak is seen at the read size of 75bp. (D) Bar plot of two ChIP-seq replicates ^17^. Relative Strand cross correlation (RSC) values, after application of the blacklist (black) or greenscreen (green) read masks relative to the no mask control (blue).

Another commonly used metric to assess the quality of a blacklist is strand cross-correlation (SCC) of all the reads in an experiment ^9^. While true protein binding sites show strand-specific enrichment towards the 5-prime ends of the reads, peaks found in input controls lack this pattern ^3,7^. When plotting the SCC at given distances between reads from opposite strands, ChIP-seq experiments have a peak at the fragment length (usually between 200 and 350 bp), while input samples have a peak at the read length (for Illumina NextSeq 500 this equals 75 bp) ^3^. Most ChIP-seq samples have a so-called ‘phantom’ peak at the read length in addition to the fragment length peak ^7^. When we measured ChIP-Seq replicate strand-cross correlation before and after masking reads using blacklist or greenscreen, the “phantom” peak at the read length position (75 bp) was effectively removed. Again, the Arabidopsis blacklist and greenscreen were equally effective at removing this artifactual signal (Figure 2C). To quantify this improvement, we computed the Relative Strand Correlation (RSC), defined as the SCC at the fragment size divided by the SCC at the read size ^3,7^. The lower the read length signal typical of inputs, the greater the relative strand correlation (RSC) of the ChIP-seq experiment. Masking ChIP-seq reads or peaks that overlap with artificial signals using either blacklist or greenscreen regions results in a similar increase in RSC (Figure 2D). The combined data suggest that the greenscreen pipeline removes artifact ultra-high signals as effectively as the Arabidopsis blacklist.

### Effect of greenscreen or blacklist filters on ChIP-seq replicate concordance

Having established greenscreen as an effective tool for artifact removal based on established metrics ^9–10^, we next investigated its effect on assessment of ChIP-seq replicate reproducibility. Correlations between peak signals are often used to evaluate the quality of biological replicates ^18^. However, highly reproducible sequencing artifacts common to all replicates distort this metric ^10^. Previous publications showed that removing artifactual signal by applying blacklist masks revealed the true correlation structure between ChIP-seq replicates ^10^. To test replicate reproducibility before and after masking we clustered on pairwise Pearson Correlation Coefficients of read signals within called peaks. We analyzed published ChIP-seq experiments, where ChIP-seq for the same proteins was conducted in different laboratories and assessed how well these samples clustered compared to biological expectation.

In particular, we employed ChIP-seq datasets for FLOWERING LOCUS D (FD), TERMINAL FLOWER 1 (TFL1), and LEAFY (LFY) ^17,19–24^. Two of the factors probed are related: TFL1 is a transcriptional co-regulator that is recruited to chromatin by the bZIP transcription factor FD ^17^, while LFY binds different genomic locations ^25^. Our expectation is therefore that the TFL1 and FD ChIP-seq replicates will cluster together more often than either does with LFY ChIP-seq replicates (Y_k=2_, Figure 3A).

**Figure 3.**
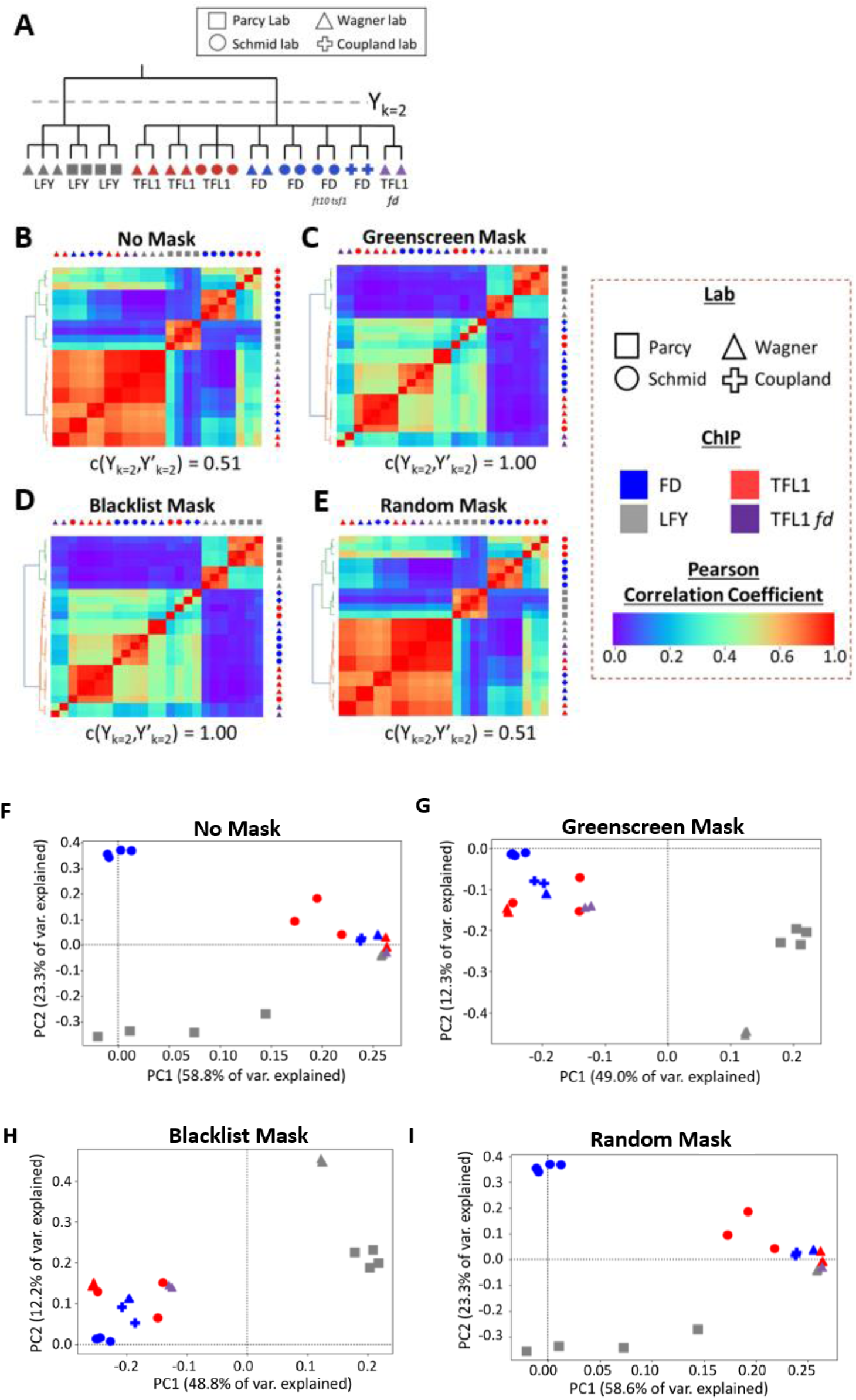
Clustering ChIP-Seq Replicates. (A) Biological expectation for clustering of 21 ChIP-seq replicates. Because TFL1 is recruited by FD, we expect all TFL1 and FD samples to cluster into one group and all LFY samples to cluster into a second group (k=2). Color: ChIP for different factors (LFY, FD and TFL1)^14–20^; Symbol: lab that conducted the ChIP. ChIP-seq peaks were called by MACS2 using published input MACS2 controls or mock MACS2 controls if input was not available. (B-E) Heatmaps of pairwise Pearson correlation coefficients. Samples are sorted using unsupervised hierarchical clustering (left of heatmaps). Legend: low (left) to high (right) correlation. Below: Rand index of clustering success relative to biological expectation. Artificial signals were either not masked (B), masked using greenscreen (C), masked using a blacklist (D), or masked using random genomic regions length-matched to greenscreen regions (E). (F-I) Principal Component Analysis (PCA) of the top two principal components. PCA were calculated using signals from the union of all ChIP-seq replicate peak sets. Artificial signals were either not masked (F), masked using greenscreen (G), masked using a blacklist (H), or masked using random genomic regions (I).

Using experiment matched inputs as MACS2 controls, or mock ChIP if no input control was available, we called peaks on non-masked reads and blacklist masked reads of each ChIP-seq replicate. In parallel, we removed peaks that overlapped with the greenscreen regions. We next calculated pairwise Pearson correlation coefficients on the replicates, performed unsupervised hierarchical clustering, and generated rand index values (Y_k=2_, Y^’^_k=2_) to measure how similar the identified clusters were to biological expectation ^26^ . Without artifact signal removal, LFY, TFL1, and FD ChIP-seq samples did not cluster according to biological expectation, yielding a low Rand index of 0.56 (Figure 3B). A similar correlation structure was observed when we computed Pearson correlation coefficients for the reads found in greenscreen regions (Figure S4). Hence the spurious correlation structure is likely due to artifact signal. However, when we applied either the blacklist read filter or the greenscreen peak mask to the ChIP-seq replicates, the expected correlation structure emerged with LFY and FD/TFL1 in separate clusters, yielding a Rand index of 1 (Figure 3C, D). By contrast, random genomic regions of similar length distribution that did not overlap with greenscreen regions did not improve the pairwise correlation coefficients nor the Rand index (Figure 3E).

The relationships between replicates can also be visualized by transforming the peak signals of each experiment from a multi-dimensional to a lower dimensional space using Principal Component (PCA) analysis to project the two principal components that make up the most variance in each ChIP-seq replicate. PCA conducted without an artifact signal mask or with a random mask only slightly separated LFY ChIP-seq replicates from the TFL1 and FD experiments in the second principal component (Figure 3F). However, after greenscreen or blacklist masking, the first principal component clearly distances the replicates or experiments by factor assayed, as expected (Figure 3G, H). Random masks were indistinguishable from no mask (Figure 3 B, I). Our combined data reveal that greenscreen is as effective as blacklisting in improving analysis of ChIP-seq replicate concordance.

Although ENCODE blacklists were generated using hundreds of inputs ^9–10^, in Arabidopsis 20 inputs sufficed to generate effectively masks for artifact reads. This prompted us to test the efficacy of greenscreen masks generated using fewer inputs. We tested three, four, six, or ten inputs randomly chosen from our twenty inputs (Table S1) for building the greenscreen mask and applied it to the ChIP-seq replicates. We found that as few as 3 inputs significantly increased rand indices for unsupervised clustering of the LFY, FD, and TFL1 ChIP-seq experiments (Figure 4A). Next, we contrasted the effect of generating a greenscreen mask using three experiment matched inputs to that derived from three unmatched inputs from Table S1. Inputs generated for a given experiment were as effective at removing artifact signals as unrelated inputs based on Rand indices (Figure 4B).

**Figure 4.**
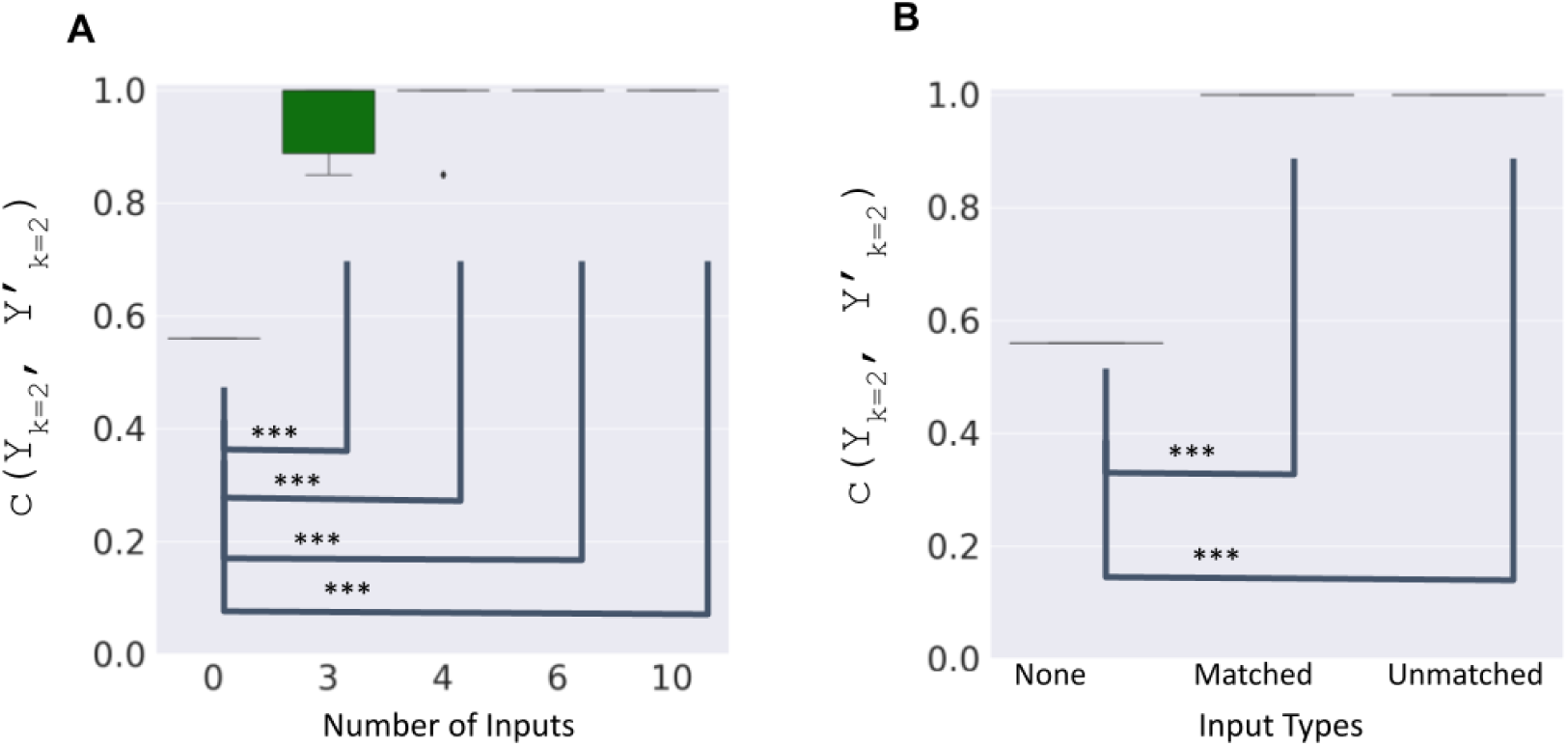
As few as three inputs effectively remove artifact signals using greenscreen masks. The distance between expected (Y_k=2;_ Figure 3A) versus true (Y’_k=2_) ChIP-seq replicate computed using rand-index values for greenscreen filters generated using numbers of input. (A) Rand-index values c(Y_k=2_, Y’_k=2_) for greenscreen filters derived from subsamples (n=3,4,6,10) of the twenty random inputs (Table S1). Subsampling was conducted 10 times with replacement. One-sample t-test (n=10) p-values relative to no mask: 3 inputs ***p = 3.4 e-08; 4 inputs: ***p= 7.3 e-9; 6 inputs ***p = 0; 10 inputs ***p = 0. (B) Rand-index values c(Y_k=2_, Y’ _k=2_) for greenscreen filters derived from five different combinations of three experiment-matched inputs ^14,16^, or of three unmatched inputs (Table S1). One-sample t-test (n=5) p-values relative to no mask: match ***p = 0; unmatched ***p = 0 (A, B) Box and whisker plots: median (center line), quartiles (box), remainder of the distribution up to 1.5xIQR (whiskers). Values past the IQR are shown as outliers.

### Greenscreen effectively masks artifact signals in larger genomes

To test the efficacy of greenscreen in a large and more repetitive genome, we next focused on human cell lines. We developed a greenscreen filter from twenty inputs selected randomly from the hundreds of inputs used for the human blacklist ^10^. We essentially followed the procedure employed for the Arabidopsis greenscreen filter (inputs using MACS2 (q-value < 10-10), peaks present in >10 inputs). However, we merged signal peaks into contiguous regions if they were less than 20 kb apart, as was done for the human blacklist ^10^, because of the larger genome size. We then applied our greenscreen filter and the published human blacklist, which was generated using 636 human inputs ^10^, to 42 ChIP-seq replicates derived from twenty ChIP-seq datasets. The mapped reads from these datasets were from nine labs and thirteen different cell lines ^11^. We performed peak calling on unmasked and blacklist masked reads and applied the greenscreen filter to the peaks called from unmasked reads.

Based on Pearson correlation analysis, the human ChIP-seq samples showed a correlation structure before masking (Figure 5A). Application of the greenscreen or the blacklist filter yielded near identical results and revealed a different Pearson correlation structure (Figure 5B, C). Importantly, the filtering correctly recovered known correlations between factors. For example, HNRNKP and PCBP2 occupy similar states of ENCODE-annotated genome segmentation that differ from those occupied by FUS ^27^. This relationship is also recovered by PCA analysis after either greenscreen or blacklist filtering (Figure 5D-F).

**Figure 5.**
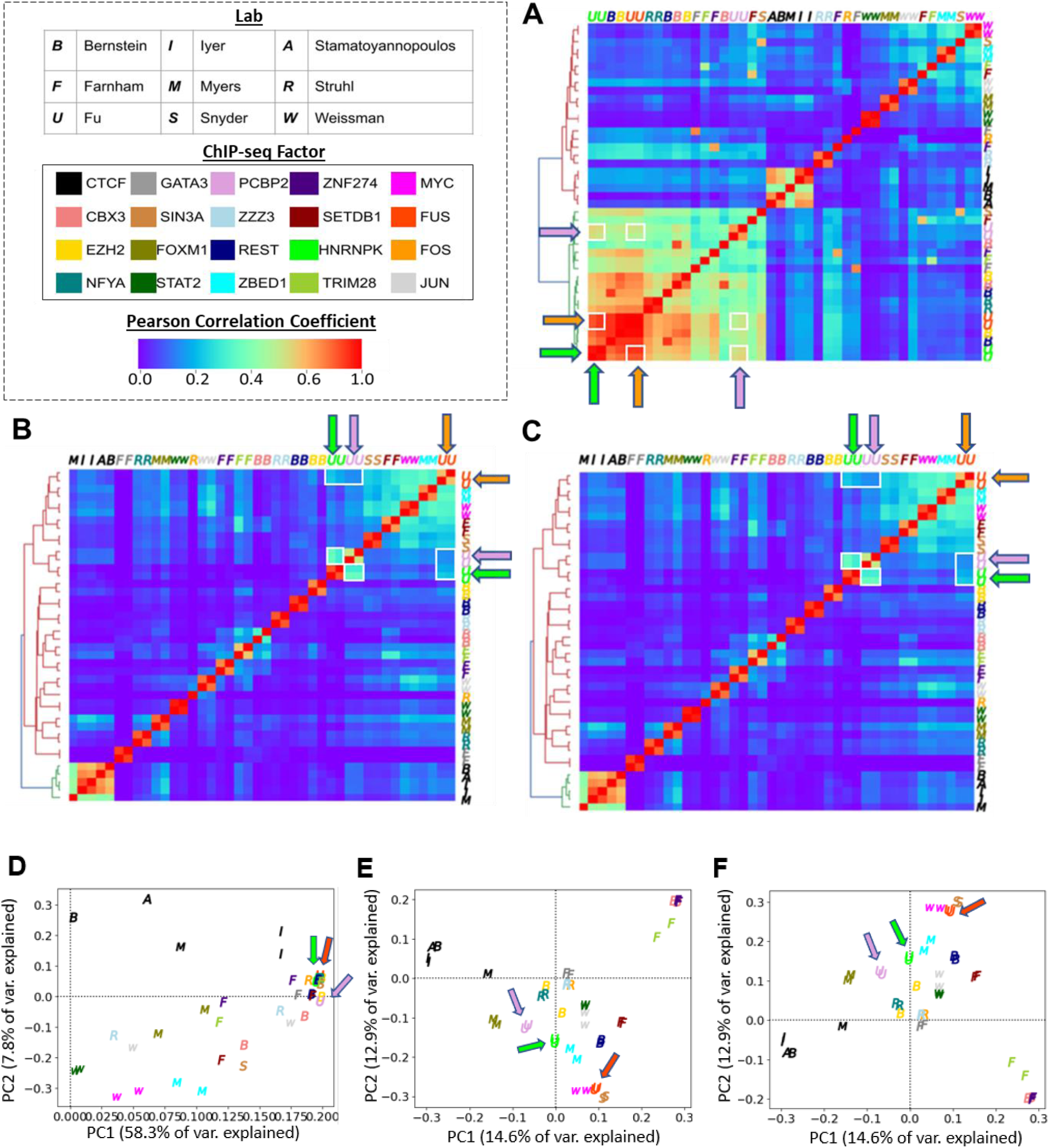
Greenscreen is as effective as blacklist in removing human artificial signal. Signal in the merged peak regions of ChIP-seq replicates from 20 different ChIP experiments conducted in 13 human cell lines by 9 labs (described in Ref ^11^). (A-C) Signals within peak regions were used to calculate and plot Pearson correlation coefficient values. Pairwise correlations between HNRNKP, PCBP2, and FUS are highlighted by white boxes and arrows. (D-F) Scatterplot of samples transformed using the top two principal components. Arrows show HNRNKP, PCBP2, and FUS samples. (A,D) no filter, (B,E) Greenscreen filter, (C,F) Blacklist filter. Color: ChIP for different factors; Letter: lab that conducted the ChIP.

Given the known genome segmentation state preference of HNRNKP, PCBP2 and FUS ^27^ (see dendrogram at top-right of Figure S5), we calculated a rand-index for unsupervised hierarchical clustering of Pearson correlation coefficients before masking (c(Y_k=2_,Y’ _k=2_) = 0.56), after blacklist (c(Y_k=2_,Y’ _k=2_) = 1.0), or greenscreen masking (c(Y_k=2_,Y’ _k=2_)=1.0). Blacklist and Greenscreen filters similarly increased the rand index using Pearson correlation analysis (Figure S5A-C). Likewise, samples transformed using PCA showed the biological relationship more clearly in the top principal component after blacklist or greenscreen masking than with no mask (Figure S5D-F).

We next used this rand-index metric to test the efficacy of human greenscreen filters generated with fewer inputs. Randomly sampling ten times groups of 10, 6, 4, and 3 different input controls we found that as few as three inputs were sufficient to generate a greenscreen filter that effectively removed artifact signal in human ChIP-seq datasets (rand-index c(Y_k=2_,Y’_k=2_)=1.0 with no variance) while only covering 0.01 % of the genome and masking 26 transcripts (Table S3). Thus, although the published human blacklist masks used over 600 inputs and masked over 227,162 kb and 3,390 transcripts ^10^ (Table 1, Table S3), greenscreen filters generated with 20 or 3 inputs were as effective as the blacklist in removing artifact signal in ChIP-seq datasets from human cell lines. We conclude that greenscreen can be applied to remove artifact signals in any new organism by using a single ChIP experiment with at least three matched inputs.

Additional methods assess factor binding to genomic locations, including Cut&Run (Cleavage Under Targets and Release Using Nuclease), which targets micrococcal nuclease to a chromatin bound factor to specifically liberate the factor associated genomic DNA ^28,29^. Cut&Run requires less tissue and is often performed without crosslinking ^28,29^. We found that Cut&Run experiments also harbor artifactual ultrahigh signal peaks and that these peaks overlap with greenscreen regions (Figure 6) ^30^. The data suggest that greenscreen can also be applied to other types of genomic datasets that probe association of a factor or modification with the chromatin.

**Figure 6.**
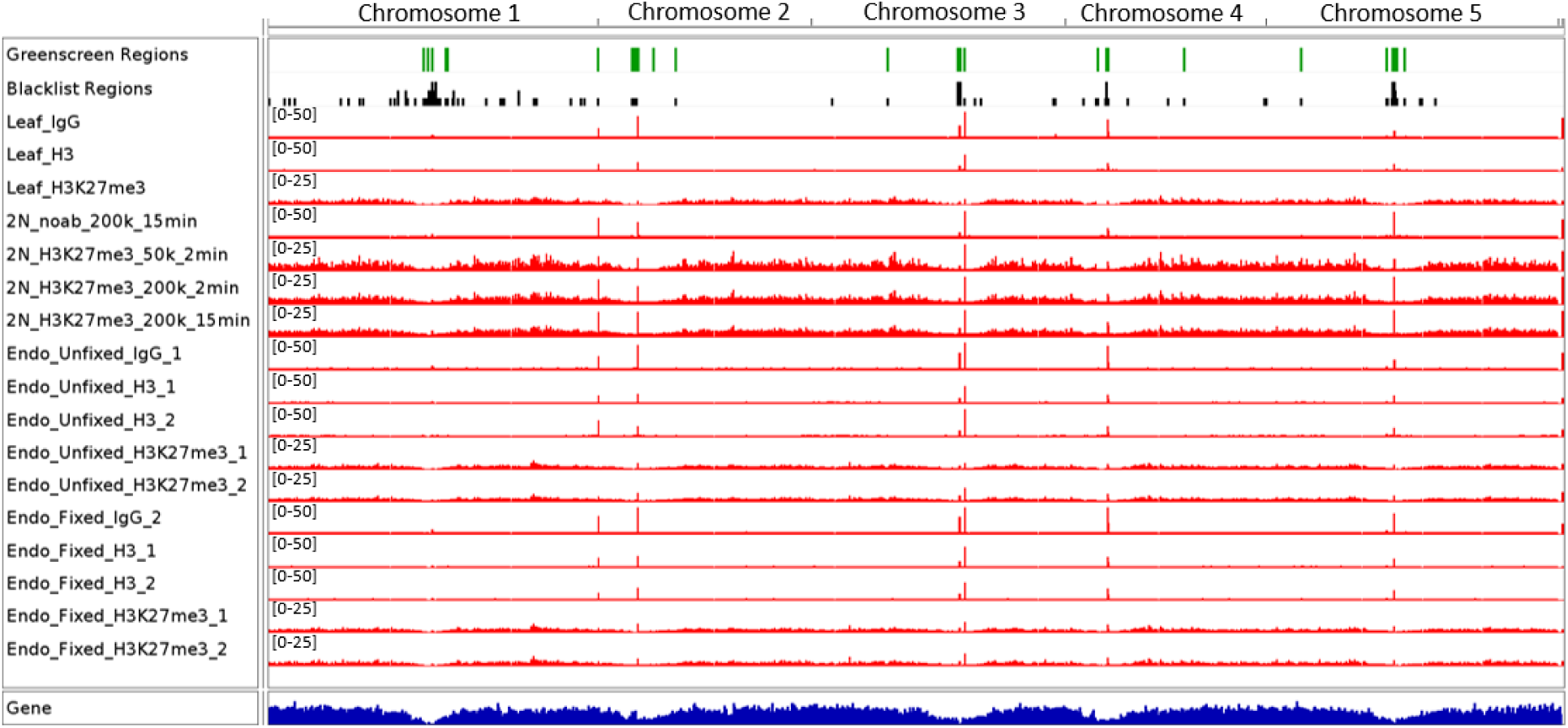
Ultrahigh signal in *Arabidopsis* Cut & Run correlates with greenscreen regions. Top: Greenscreen and blacklist regions. Below: Cut&Run bedgraph files for IgG, H3 and H3K27me3 Cut&Run from fixed and non-fixed tissues accumulation in Arabidopsis endosperm and leaves ^30^. Ultrahigh artifact peaks overlap with greenscreen regions.

### An improved ChIP-seq analysis pipeline that incorporates greenscreen

We next assessed the effect of greenscreen and of different commonly used ChIP-seq analysis parameters on peak calling in the above-mentioned LFY and FD ChIP-seq datasets. Towards this end, we defined potential true positive peaks as peaks present in both a LFY ChIP-seq dataset ^20^ and an orthogonal LFY ChIP-chip dataset ^25^, but do not overlap the union of blacklist and greenscreen regions. For FD we compared two FD ChIP-seq experiments from different laboratories ^17,19^ in an analogous manner. We labelled ChIP-seq peaks that overlapped with the union of all greenscreen or blacklist regions as probable false positive peaks, or artifacts.

For ChIP-seq analysis, we merged LFY or FD ChIP peak signals from replicates in MACS2 after down sampling to achieve equal genome coverage (see methods for detail) and identified significant (MACS2 summit qval≤10^-10^ cutoff) peaks using either no control, input, or mock controls. Analysis without controls yielded a large number of potentially ‘false peaks’ that overlapped with the combined greenscreen and blacklist filter (Figure 7A, Figure S6A). By contrast, input controls removed some, and mock controls eliminated all probable false positive peaks in both the LFY and the FD ChIP-seq datasets (Figure 7A, Figure S6A). However, relative to input normalization, mock controls also removed 22% (LFY) or 12% (FD) of the likely true ChIP peaks, potentially increasing the false negative rate.

**Figure 7.**
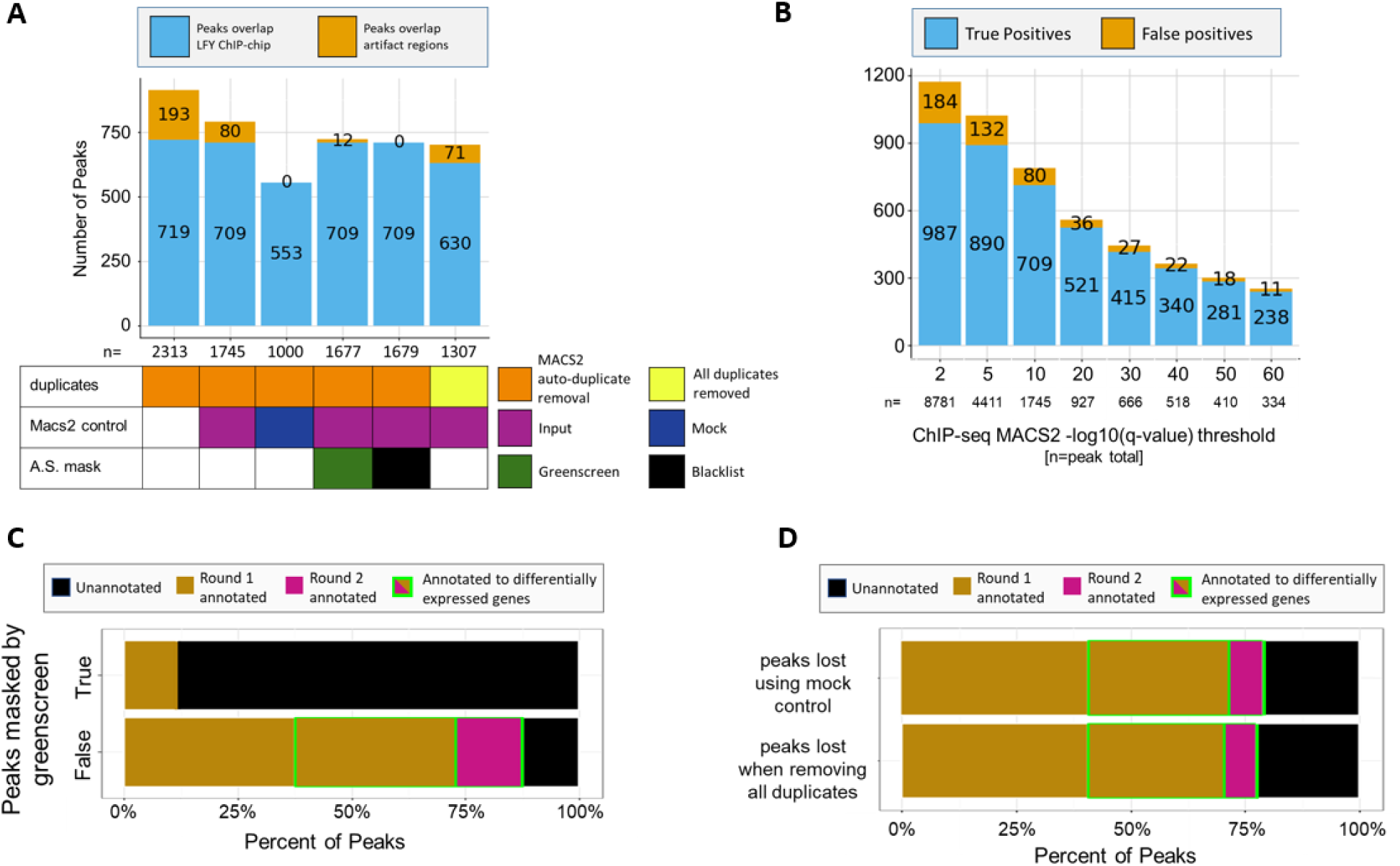
Optimizing ChIP-seq peak calling pipeline with filtering. (A). Stacked bar chart to assess the impact of calling LFY ChIP-Seq MACS2 peaks using no control, input or mock control; duplicate removal (MACS2 keep dup auto or no duplicates) and masking (none, greenscreen and blacklist). Total peak number (“n” value under x-axis), potential true positive peaks (LFY ChIP-seq peaks^20^ that overlap with LFY ChIP-chip peaks^25^ ; turquoise bars with peak number) and potential false positive peaks, (peaks overlap with the union of blacklist and greenscreen regions; orange bars with peak number). (B) Stacked bar chart to assess the impact of increasing the ChIP-seq peak summit q-value threshold for calling LFY ChIP-seq peaks. (C) Horizontal stacked bar charts for ChIP-seq peaks (MACS2 ‘keep dups auto’ using input controls), Top: ChIP-seq peaks that overlap with greenscreen (n=68). Bottom: ChIP-seq peaks do not overlap greenscreen (n=1677). Peaks were assigned to genes (brown and pink) as previously described ^20^ or could not be assigned (black), we also indicate whether genes were rapidly differentially expressed in response to factor binding (green box) ^20,25^. (D) Horizontal stacked bar charts for LFY ChIP-seq peaks analyzed and in (C) that were lost when using mock instead of input control in MACS2 (n=693; top) or when removing all duplicates instead of using MACS2 ‘keep dups auto’ (n=456; bottom).

Since input controls generally increased MACS2 peak calling specificity, we next tested the effect of duplicate removal, artificial signal masking, and summit q-value thresholds in input normalized ChIP datasets. It is quite common to remove all duplicates from ChIP-seq data analysis even though a more nuanced approach was proposed as duplicate reads are known to contribute to true ChIP-seq signal ^5–6,9,31^. One such approach is the MACS2 default is ‘keep dups auto’, which removes duplicates in excess of expectation based on the effective genome length and sampling depth, and those that do not fit a binomial distribution at a given location (p≤1e-5) ^13^. We found that removing all duplicates lowered, but did not eliminate, peaks overlapping with artifact signals and simultaneously reduced the number of likely true peaks (Figure 7A, Figure S6A). As expected, the greenscreen or blacklist filters removed the majority of peaks that overlapped with universal artifacts, and neither approach adversely affected true peak number (Figure 7A, Figure S6A). A small number of ChIP-seq peaks were in blacklist but not in greenscreen regions, and hence remained after applying the greenscreen filter. However, these regions were characterized by low input (artifact) signal and are likely evidence of over masking by the blacklist (Figure S7). Finally, increasing peak calling stringency by decreasing the minimum summit q-value threshold was not an effective means to remove artifact peaks and strongly reduced the final true peak set (Figure 7B, Figure S6B). This is expected; artifact peaks contain ultra-high signal and thus have highly significant summit q-values.

To further test the validity of our peak designations, we examined the properties of the 1677 LFY and 5294 FD “true” ChIP-seq peaks called by applying our ChIP-seq pipeline (replicate down sampling before merge, ‘keep dups auto’ in MACS2, input controls and greenscreen filter) and of the “false” ChIP peaks removed by the greenscreen mask. We assigned all peaks to genes as previously described ^17,20^ (see methods). Most of the LFY and FD peaks identified using the new ChIP-seq pipeline were located near protein coding or microRNA genes. Furthermore, more than half or one quarter of these genes, respectively, were rapidly differentially expressed after LFY or FD activation (Figure 7C, Figure S6C). By contrast, the majority of the peaks removed by greenscreen in both LFY and FD ChIP were not located near genes nor differentially expressed. This suggests that the peaks removed by greenscreen are false positives. Indeed, the ChIP-seq peaks which overlapped greenscreen regions had SCC signal peaks at the read length (Figure S8), like the artifactual signal in input ^7^.

The distinct properties of the true and false ChIP-seq peaks prompted us to examine the peaks lost when using mock rather than input control in MACS2 (693 for LFY and 1650 for FD) and the peaks lost when removing all duplicates rather than MACS ‘keep dups auto’ (456 for LFY and 1460 for FD; Figures 7A and S6A). We found that the peaks removed in either case mapped near genes and showed differential expression at similar levels as true ChIP-seq peaks (Figure 7D, Figure S6D). Thus, mock normalization and all duplicate removal eliminate true positive peaks. In conclusion, calling peaks using MACS2 duplicate presets and input controls, followed by the greenscreen filter, maximizes the potential true positive peak number while effectively removing artifacts.

### The improved ChIP-seq pipeline, which incorporates greenscreen, identifies more true peaks

To test the ChIP-seq pipeline (replicate down sampling before merge, ‘keep dups auto’ in MACS2, input controls and greenscreen filter) we applied it to ChIP-seq datasets described in this paper^17,19–24^. Relative to published datasets ^17,19–24^, our pipeline generally identified at least twice as many peaks (figure margins; Figure 8A) To assess whether these newly identified peaks are true peaks, we performed a pairwise peak overlap analysis. We computed the fraction of peaks in a given experiment (rows in Figure 8) that overlap ChIP-seq peaks identified in a second experiment (columns). For pairwise comparisons using the same binding factor, peak overlap was consistently higher with ChIP-seq peaks identified using the new analysis pipeline. Hypergeometric tests revealed equal significance for the peak overlaps of the new pipeline and the published (smaller) peak sets (Figure S9). Thus, the new ChIP-seq pipeline, which includes greenscreen masking, identified more true positive peaks than published procedures. Moreover, the newly identified target genes are differentially expressed rates similar to those identified by the published approaches (Figure S10). In addition, the new ChIP-seq pipeline identified genes in pathways previously linked to FD/TFL1 ^17^ in several of the datasets analyzed, including the chromatin regulators PICKLE RELATED 1 (PKR1) and BRAHMA (BRM), the strigolactone hormone response factor SUPPRESSOR of MAX2 LIKE 6 (SMAXL6) and the meristem identity regulators LATE MERISTEM IDENTITY 2 (LMI2) and FRUITFULL (FUL) (Figure 8B). For LFY several new target genes were gained in all datasets, such as the transcription factor PEAR1 which controls vascular development and the auxin conjugating protein DWARF in LIGHT1 (DFL1), while in other cases new targets are found in only some of datasets (chromatin regulator JMJ30)^32–34^ (Figure 8C). Thus the improved ChIP-seq pipeline removes artifact signal, calls more true peaks, and identifies additional biologically relevant target genes.

**Figure 8.**
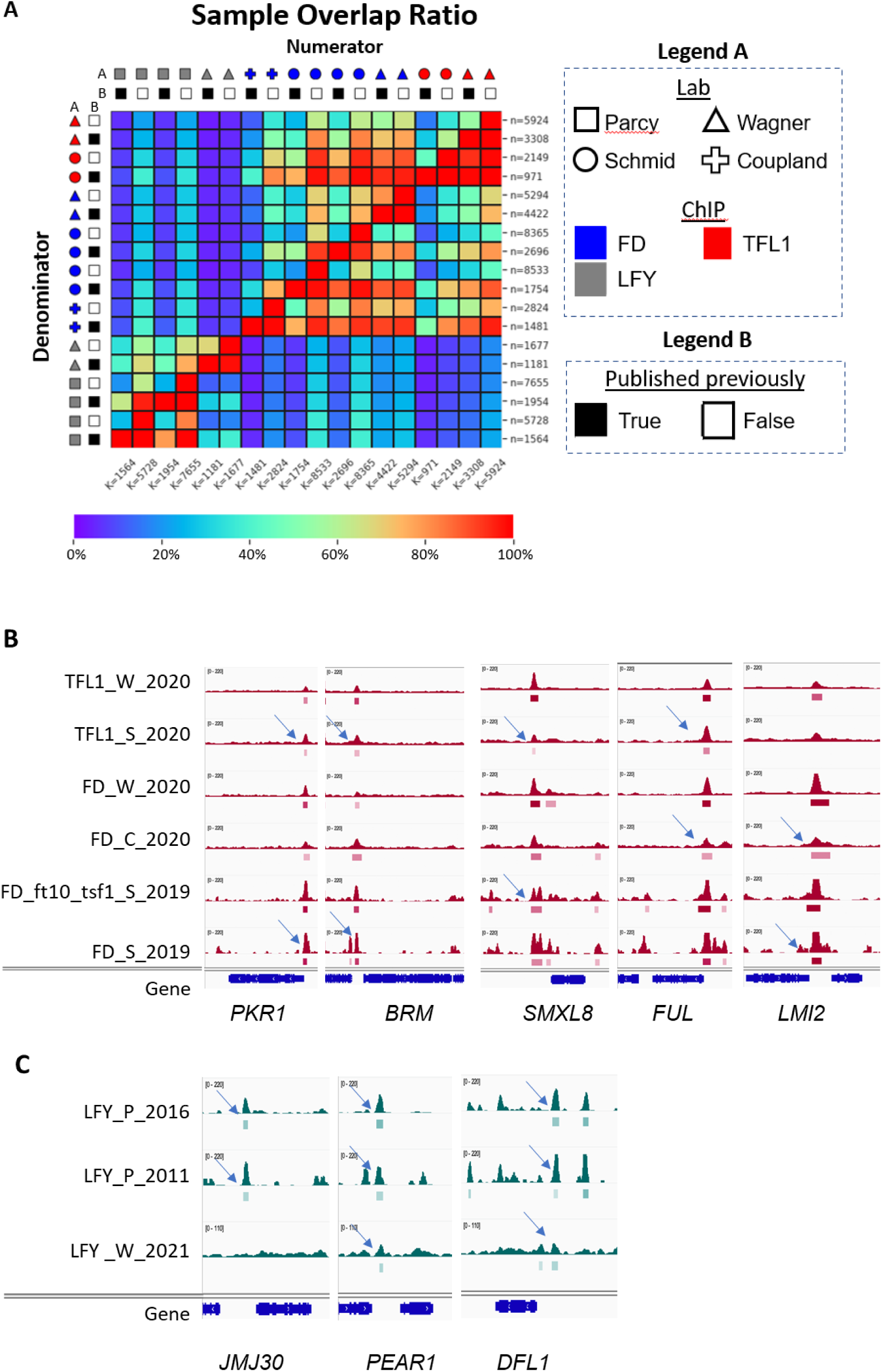
Improved ChIP-Seq pipeline calls many more true peaks than published methods. Heatmap for the ratio of peaks from experiment A in each row that overlap with the peaks from experiment B in each column in pairwise comparisons divided by the total peaks in experiment A. The total number of peaks per experiment are listed on the perimeter of the heatmap. ChIP-seq samples of three factors (LFY, FD and TFL1), conducted in four different laboratories ^17,19–24^ were analyzed (Legend A). MACS2 controls were matched to corresponding publications. Peaks were either identified using the optimized ChIP-seq method (published previously=False) or using the methods published (published previously=True) (Legend B) Note that published datasets from the Wagner lab were already greenscreen filtered ^17,20^. Legend: percent overlap. Raw numbers for the heatmap are in Table S4. (B-C) Examples of new peaks identified (arrows) by applying the optimized ChIP-seq pipeline to previously published TFL1, FD (B), and LFY (C) datasets ^17,19–24^. Browser view of ChIP-seq signals (RPK10M; all scales show range 0-220 except LFY_W_2021 (range 0-110)) with significant peaks (summit *Q* < 10^−10^) according to MACS2 marked by horizontal bars, with the saturation proportional to the summit *Q* value (as for the narrowPeak file format in ENCODE).

### Greenscreen filtering improves detection of factor binding site overlap and of factor occupancy changes

Artifact signal removal improves estimates of factor binding overlap in ChIP-seq datasets. For example, when considering binding peak overlap in the LFY, FD and TFL1 ChIP-seq datasets we analyzed, LFY and FD apparently occupy similar genomic regions in some experiments without masking or after applying a random mask based on Pearson correlation and PCA analyses (Figure 9A, D, E, H). Application of the greenscreen (or the blacklist) filter clearly separates the LFY from the FD/TFL1 bound regions by Pearson correlation and in the first principal component of the PCA analysis (Figure 9 B, C, F, G). In addition, application of the greenscreen filter allows detection of factor binding in different conditions after greenscreen filtering. LFY ChIP-seq conducted in root explants clearly separates from that conducted in reproductive tissues (Figure 9F). Thus, removal of artifactual signal by greenscreen is critical to derive biological relevant information from comparative ChIP-seq binding analyses.

**Figure 9.**
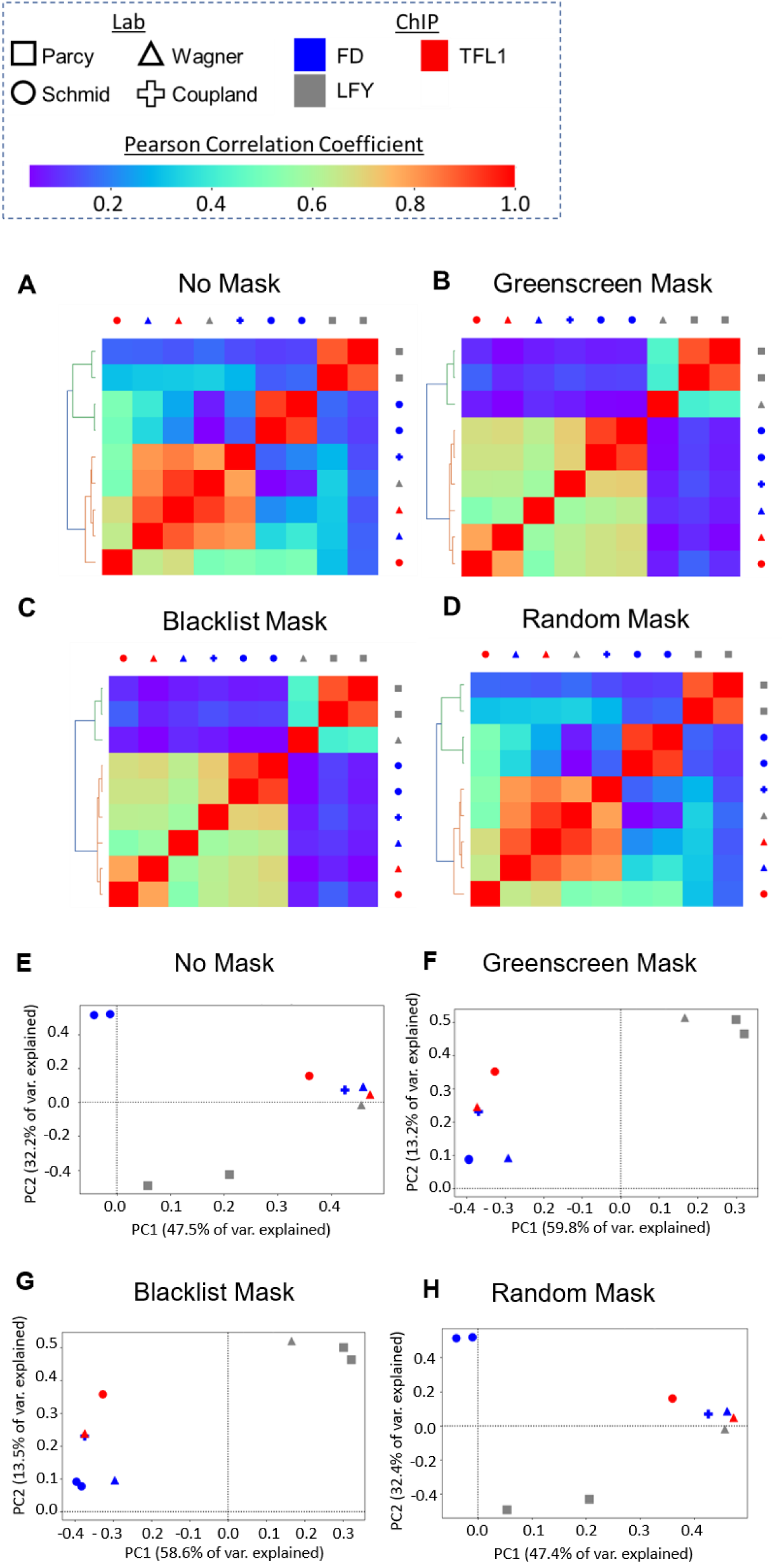
Artificial signal masks reveal biologically relevant relationships between Arabidopsis ChIP-seq datasets. Pearson correlation coefficient values for FD, LFY, and TFL1 ChIP experiments ^17,19–24^ given averaged replicates (see methods for details). Input controls were used, where none were available mock controls were employed. Above: legend for color codes and symbols. (A-D) Heatmaps organized by unsupervised hierarchical clustering of the samples based on the Pearson correlation coefficients. (A) Without an artificial signal mask, two FD and LFY ChIP-seq experiments cluster together. (B,C) After greenscreen peak masking, samples cluster by factor as quantified by the Rand-index. The same result is obtained with the blacklist. (D) A random peak mask (negative control) produced clustering similar to no mask. (E-H) Principal Component Analysis (PCA) plots of the top two principal components. Artificial signals were either not masked (E), masked using greenscreen generated using 20 inputs (F), masked using a blacklist generated with the same 20 inputs (G), or masked using random genomic regions length-matched to greenscreen regions (H).

The importance of masking on biological conclusions derived from ChIP-seq datasets is further underscored by analysis of enhancer co-occupancy of different transcription factors. A database for Human and Arabidopsis ChIP-seq and DNA affinity purification sequencing (DAP-seq) datasets called ReMap 2020 includes binding information for 372 Arabidopsis transcriptional regulators (179 ChIP-Seq and 330 DAP-seq datasets) ^35^. This catalog can be used to identify enhancers bound by many transcription factors, possible stretch or super enhancers ^35^. However, artificial signal masking was not performed, hence it is difficult to distinguish genomic regions where many proteins bind from regions of high artifact signal. Indeed, we found that 6,664 peaks and 91 “cis-regulatory modules” (CRMs) identified by Remap 2020 overlap with greenscreen regions (Figure 10). 17 of the CRMs show one-hundred or more different transcription factor binding sites. Hence, analysis of factor binding and specific identification of hotspots should include artifact signal filters.

**Figure 10.**
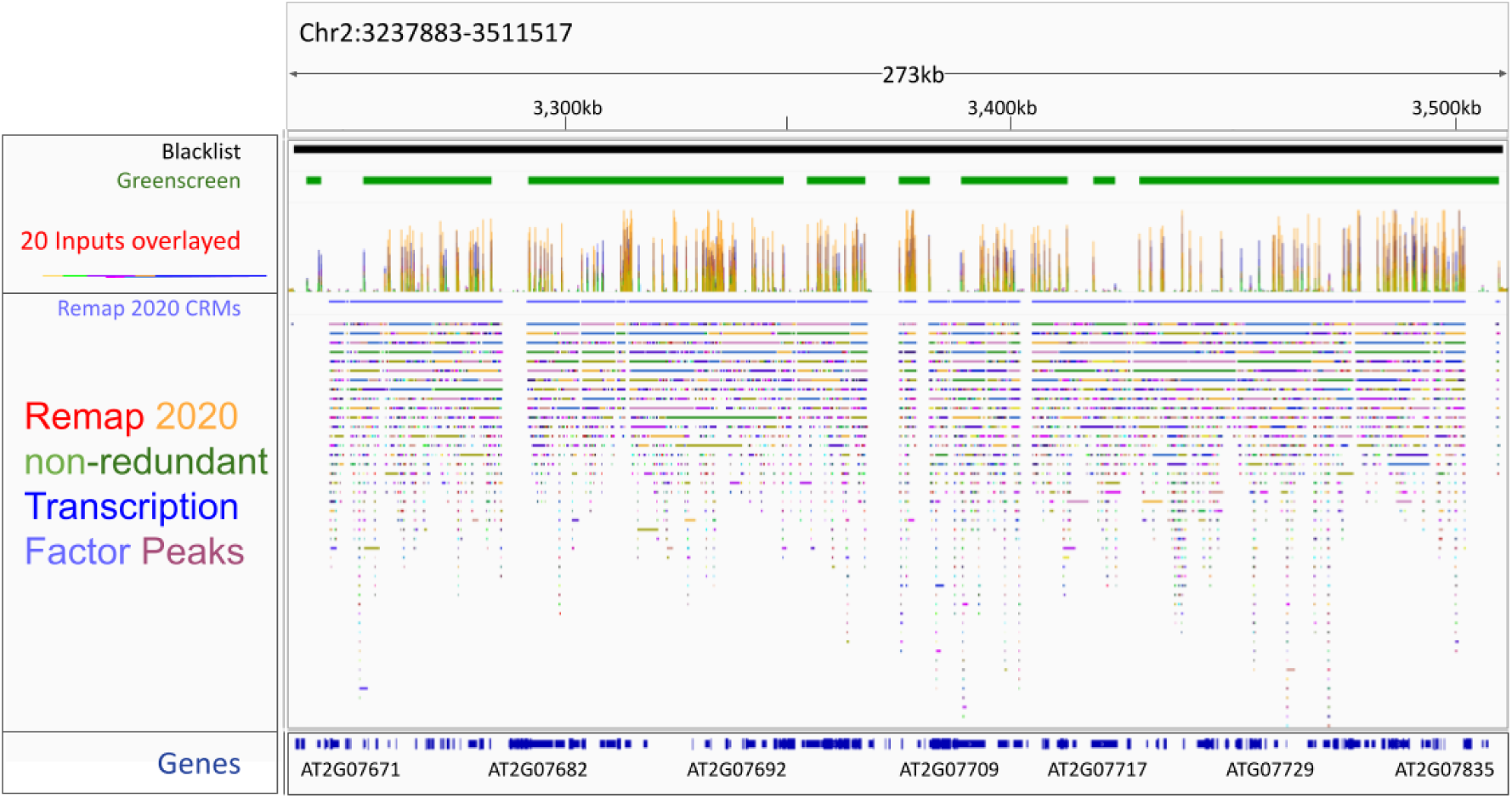
Transcription factor binding hotspots overlap with artifact signal. Genome browser view of Arabidopsis Chr2: 3,237,883-3,511,517. Header shows blacklist (black) and greenscreen (green) artifact regions (top), signals from the 20 inputs used to generate the greenscreen filer (below) and ChIP-seq factor binding hotspots called cis-regulatory modules (CRMs, blue lines) defined according to Remap 2020 as merged regions of all non-redundant peaks ^35^. Tracks: non-redundant Remap 2020 binding regions or the average start and stop sites of overlapping target sites for each given transcription factor ^35^. Different colors represent different transcription factors ^35^. Bottom: Araport 11 Gene models^49^.

Finally, masking artifact signals enables proper detection of changes in factor binding, for example in different genetic backgrounds. In Arabidopsis, PRC2 is recruited to target loci by class I BPC and C1-2iD Zn-finger transcription factors in Arabidopsis ^36^. Without greenscreen filter, depletion of PRC2 recruiting factors appears to have subtle consequences on PRC2 occupancy (Figure 11). This may be due to shared ultrahigh artifactual signal between the two ChIP-seq datasets. Indeed, application of the greenscreen filter dramatically reduced background noise and revealed the true contribution of the PRC2 recruiting factors to PRC2 occupancy. In summary employing the greenscreen filter improves detection of ChIP-seq peak overlap and allows accurate detection of factor occupancy changes in different conditions.

**Figure 11.**
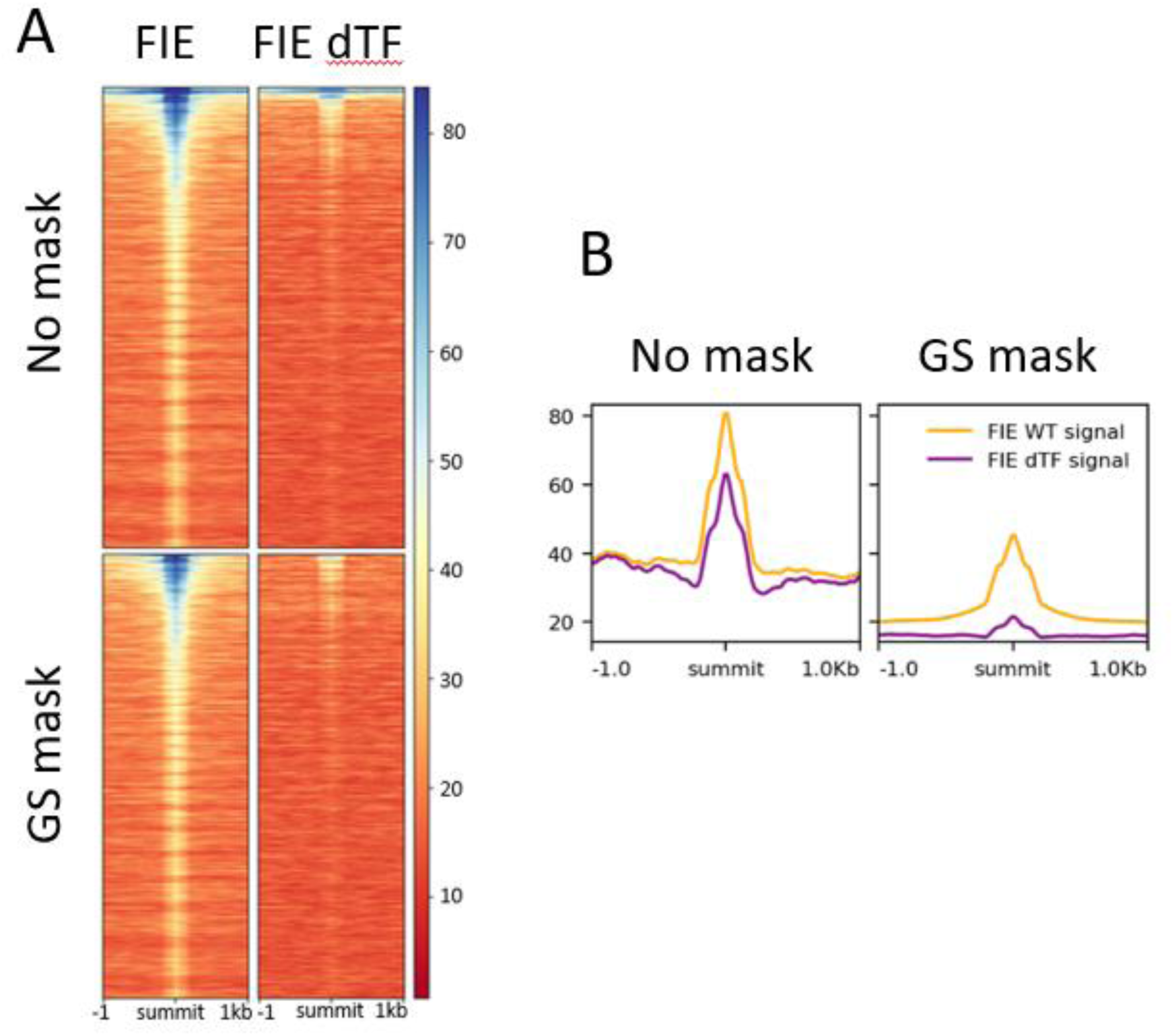
ChIP for the PRC2 component FIE in control plants and in plants that lack PRC2 recruiting transcription factors. A. ChIP-seq signal in a 2 kb region flanking significant FIE peak summits (q value ≤ 10−10, MACS2) and ranked by increasing MACS2 summit q-values for FIE without green screening (top heatmaps) and with green screening (bottom heatmaps). ChIP was performed in the wild type (FIE; left) or in plants that lack PRC2 recruiting transcription factors (FIE dTF; right) ^36^. Note the ultra-high (blue) signal at the top of the heatmaps generated with no filter. B. Mean signal of significant FIE ChIP-seq (RP10M) +/-1 kb of significant FIE peak summits without mask (right) and after applying the greenscreen mask (left). Shown are FIE in the wild type (yellow) or in plants depleted of PRC2 recruiting transcription factors (purple).

## Discussion

Artificial signal found in ChIP-seq experiments obscures true correlation between replicates or experiments, estimates of factor binding changes in different genetic backgrounds or tissues, and identification of bona fide multi-factor high-occupancy regions. The underlying mechanism by which ChIP-seq artifacts arise remains unknown, they are likely caused by multiple factors and depend on the quality of the genome annotation ^8–10^. ChIP artifacts have previously been described in many species ^2–10^ and reads are commonly masked from downstream analysis in human, mouse, worm, and fly using curated blacklists ^7–10^. However, in other model and non-model organisms, identification and removal of the artifact regions has not been standardized. Here we developed a method to identify and filter out these artifacts using a small number of inputs and tools commonly used for ChIP-seq analysis.

As for blacklists, greenscreen parameters should be optimized based on both the size of the genome and the quality of the genome build. Of particular importance was the ultrahigh signal merge parameter. For *Arabidopsis,* which has a compact genome, a 5 kb merge was optimal (Table S2). Independently, for *Drosophila*, which has a genome size comparable to *Arabidopsis,* the ENCODE Drosophila blacklist chose a 5kb merge parameter ^10^. For humans, we employed a 20 kb merge, in accordance with the ENCODE blacklist ^10^. By contrast, input peak calling (MACS2 <=10^-10^) and number of inputs needed to build greenscreen filter were similar for Arabidopsis and human.

By identifying and masking ChIP-seq peaks with the greenscreen filter, we improved enrichment of true-positive peaks and decreased erroneous correlations, revealing true biological signals. Application of greenscreen masking to Arabidopsis ChIP-Seq experiments from different labs was as effective at removal of artifact signal as the Arabidopsis blacklist, on the basis of metrics commonly employed to assess ultrahigh artifact signal removal (SSD, RSC, SCC) ^9,11^. Both filters performed equally well in revealing true ChIP-seq replicate and experiment concordance using Pearson correlation or principal component analyses in Arabidopsis and human ChIP-seq data. In both Arabidopsis and human ChIP-seq datasets, greenscreen masks a smaller percentage of the genome (0.41% and 0.01 %, respectively) compared to blacklisting. The biggest difference between blacklist and greenscreen is that the blacklist software identifies broader regions. Additional experiments are needed to determine whether the blacklist software is over masking the genome.

In non-model or new model genomes, few sequenced inputs are generally available. In *Arabidopsis* greenscreen filters generated from twenty inputs performed well, moreover, a greenscreen filter generated from 20 inputs generated a remarkably similar correlation structure for human ChIP-seq replicates as the published blacklist, which is based on over 600 inputs ^10^. Indeed, based on Rand indices, as few as three inputs are effective at artifact signal removal in both Arabidopsis and human ChIP-seq datasets, indicating that greenscreen can be used to improve ChIP-seq data analysis in any new species. The inputs can be from the same experiment as the ChIP-seq datasets as matched and unmatched inputs performed equally well. An additional advantage of the small number of inputs required is that it is easy to generate a new greenscreen filter if a new reference genome build is released, or in conditions where massive genome re-arrangements occur, such as in cancer cell lines ^37,38^. This flexibility is also useful for partially assembled genomes or polypoid genomes, like those of many crops.

We present an improved ChIP-seq analysis pipeline that incorporates greenscreen. This pipeline uses replicates with equal genome overage, input controls and the MCAS2 preset ‘keep dups auto’. We employ down sampling of high-quality replicates to equal genome coverage to avoid the replicate with the highest sequencing depth from dominating analyses in MACS2. Other approaches have been proposed to address this issue ^39^ and future efforts are needed to improve replicate handling in ChIP-seq analyses. In addition, while mock controls can remove artifactual signal, they also obscure true ChIP-seq peaks. Because of this and their low overall signal and high variability, mock ChIP is not a suitable control for ChIP-seq peak calling ^40^. Finally, we find that removing all duplicates does not effectively remove artifact signal and leads to removal of true ChIP-seq peaks, in agreement with prior studies showing that duplicate reads contribute to ChIP-seq signal ^5–6,9,26^. The improved ChIP-seq pipeline, which includes the greenscreen filter, calls more significant peaks compared to published analyses. The additional peaks show strong overlap with other ChIP-seq datasets for the same factor and link to functionally important, differentially expressed, genes and pathways.

Finally, we show that removal of artifactual signal with a greenscreen filter increases one’s capacity to form accurate biological conclusions from factor binding or chromatin modification genomic datasets, such as ChIP-seq and likely also Cut&Run. On one hand, artifact removal allows identification of precise overlap between factor binding or chromatin modification locations and true binding hotspots. On the other, it allows detection of true factor occupancy changes in different conditions.

In summary, our ChIP-seq pipeline, which incorporates greenscreen, shows high sensitivity for true peak detection. Greenscreen results in fewer artifact signal regions that are narrower than blacklist regions, and therefore removes fewer peaks. Yet both methods improve downstream analysis equally. Moreover, greenscreen filters are generated with common ChIP-seq analyses tools and using very few inputs and hence can readily be adapted to any new organism or genome.

## Materials and Methods

### Identification of artifact signal in ChIP-seq

Single-end reads from twenty ChIP-seq input controls in *Arabidopsis* were retrieved from different experiments (Table S1). FASTQC (v0.11.5) ^41^ was used to assess the quality of each sample. Inputs were not considered for downstream analysis if the average reads did not have sequencing qualities above Phred33 score 30. After passing the sequencing quality criteria, inputs were cleaned with Trimmomatic v0.39 (LEADING:3 TRAILING:3 SLIDINGWINDOW:4:15 MINLEN:36) ^42^. If needed, the remaining adaptor sequence was removed using the trimmomatic ILLUMINACLIP function (2:30:10). Trimmed reads were then mapped with bowtie2 v2.4.1^43^ to the TAIR10 ^14^ *Arabidopsis* genome using default parameters. Reads that did not map, did not generate a primary alignment, did not pass quality checks, did not map to a nuclear chromosome or had MAPQ≥30 was eliminated from downstream analyses.

To ensure that the samples were, in fact, ChIP-seq input controls we generated SCC plots using MACS2 predictd function ^13^ for each input and removed any sample that returned a ChIP-seq experiment signature from a Watson and Crick strand correlation test (Figure S11). SCC metrics are typically used for quality control of ChIP-Seq to quantify an experiment’s signal-to-noise ratio ^9,11^. In ChIP-Seq, distinct Watson and Crick strand read enrichment occurs on opposite sides of a factor binding site at least a DNA fragment length apart. Since input does not include immunoprecipitation of factor bound DNA sequences, input should not show enrichment at the fragment size or above on a strand-cross correlation plot ^9^. We used both MACS2 v2.2.7.1 ^13^ and ChIPQC 1.26.0. ^16^ to assess input quality .

### Blacklist generation

Blacklist generation requires both uniquely mappable regions in the genome and mapped input reads (Figure S2). Uniquely mappable sites were annotated using UMap (1.1.0) ^12^ on the TAIR10 *Arabidopsis thaliana* genome assembly ^14^. In parallel, high quality mapped reads (MAPQ >=30) from twenty input samples were retained for import into the blacklist tool and employed in the greenscreen pipeline (below). The blacklist tool is hard-coded for blacklist regions in the human genome. To account for the smaller genome size of *Arabidopsis* we manually modified the code (blacklist.cpp, line 469) to merge regions within 5kb rather than 20kb.

### Greenscreen generation

Utilizing the same mapped reads used to generate the blacklist, we identified peaks from each input sample individually using MACS2 v2.2.7.1 ^13^ (—keepdup “auto” —no model –extsize [read length] —broad --nolambda -g 101274395). By default, MACS2 identifies significant signals with a dynamic Poisson distribution by capturing the local backgrounds in its lambda parameter ^13^. We set ‘--nolambdà to ensure we capture ultra-high signals above the global background specifically. Additionally, bypassing the default MACS2 shifting model which extends reads based on what MACS2 estimates to be the samples fragment length, reads were extended in 5’-3’ direction based on each sample’s read length as determined by ChIPQC 1.26.0 ^16^. The effective genome size was fixed to 85% of the full *Arabidopsis* genome size ^44^.

To optimize the greenscreen mask we strove to minimize false-positive ChIP peaks called, while also minimizing the percent of the genome and the number of genes masked (Table S2). ChIP-seq improvement was measured as enrichment of peak overlap between ChIP-seq and ChIP-chip datasets for the same transcription factor (Table S2). In addition, we quantified expected unsupervised clustering of pairwise Pearson correlations between ChIP-seq replicates for the same factors from different laboratories by calculating rand-index values (see Figure 3). These combined investigations defined optimal greenscreen artifact peaks as those with an MACS2 q-value < 10^-10^ and merging of peaks with maximum merge distance of 5 kb. After removing peaks with q-value >=10^-10^ (column 9 in the broadPeak output file) from each of the twenty inputs, we concatenated all input peak regions. Lastly, we removed those regions that did not have significant artifact peaks in at least half of the inputs analyzed.

### ChIP-Seq peak calling

Raw Arabidopsis read data was obtained and cleaned using Trimmomatic v0.39 (LEADING:3 TRAILING:3 SLIDINGWINDOW:4:15 ILLUMINACLIP:TruSeq3-SE.fa:2:30:10 MINLEN:36). After trimming, reads were mapped (MAPQ >=30) to the TAIR10 *Arabidopsis* genome using default parameters in bowtie2 v2.4.1. To implement blacklist masks, the reads in ChIP-seq and controls samples that overlapped blacklist regions were removed from downstream analysis using samtools v1.7 (htslib v1.7) ^45^.

Next, MACS2 v2.2.7.1 ^13^ (—keepdup “auto” —nomodel –extsize [fragment_length] -g 101274395) was utilized to perform peak-calling on the ChIP-seq samples with and without the blacklist mask^13^. Reads were extended within the MACS2 software in 5’-3’ direction based on each sample’s fragment size, determined by ChIPQC 1.26.0 ^16^. To balance removing duplicates generated from PCR amplification verses duplicate reads which originate from independent fragment. it is recommended to not eliminate all duplicates but instead set a duplicate threshold per genomic location based on an experiment’s sequencing depth^5–6,9,31^. Therefore, duplicate reads were managed in MACS2 v2.2.7.1 by setting “--keepdups auto”. To pool ChIP-seq and controls replicates, reads were randomly down-sampled using biostar145820 ^46^ to match the read-depth of the replicate with the lowest read depth. This requires high quality replicates; we recommend only using replicates that have peak signals with Pearson correlation co-efficient > 0.7 after applying the greenscreen filter. Additional ChIP-seq quality measures were discussed ^47^. Throughout, we conducted MACS2 using experiment-matched normalization controls by scaling the ChIP-seq signal to input when available, otherwise mock control (Table S3). Reads with a summit q-value (column 9 in MACS2 narrowPeak output) less than or equal to 10^-10^ were retained. On peak calling files generated without blacklist mask, we performed greenscreen using bedtools (v2.26.0) ^48^ to remove peaks that overlapped with greenscreen regions.

To annotate peaks to genes, we used Araport11 gene annotation of the *Arabidopsis* genome ^49^. Peaks with intragenic summits were first annotated to the genes to which they were intrinsic. Remaining peaks with summits at least 3kb upstream of a gene were then annotated to the closest upstream peak. If data was available for rapid (immediate early) gene expression changes after factor binding, we mapped the remaining orphan peaks within 10kb (upstream or downstream) of significantly differentially genes (round 2 annotation)^17^.^20^. Annotation code is available on github: https://github.com/sklasfeld/ChIP_Annotation.

### ChIP-seq Pearson Correlation and PCA plots

Pearson correlation and PCA plots assess the signal within regions of interest. All of the ChIP peaks called in each replicate (Figure 3) or pooled samples (Figure 9) were concatenated and merged to a final bed file to be used as regions of interest. Signal files were generated after read extension to the respective sample’s fragment size and normalization over all mapped ChIP-seq reads or over all reads left after blacklist masking. Blacklist masking requires read removal, while greenscreen relies on peak removal.

The normalized signal within each of the selected regions was arranged into a matrix. Pairwise Pearson’s correlation was calculated across the columns, and then unsupervised hierarchical clusters (k=2) were generated using Ward’s clustering methods ^50,51^. Heatmaps and dendrograms were plotted to display these results. To quantify how well the unsupervised clusters match our hypothesis of how they should cluster based on the known biology we calculated the rand-index ^26^. A tutorial on how to run the pipeline to generate these results can be found at: https://github.com/sklasfeld/GreenscreenProject.

Similarly, to visualize the samples using the top two principal components, the signals with each of the sample’s peak regions were also measured using deeptools (3.5.1) multiBigwigSummary ^52^. Values for each sample in the top two principal components were plotted with deeptools (3.5.1) plotPCA function^52^.

### Human greenscreen filter generation

To identify inputs suitable for greenscreen analysis, we applied ChIPQC to the hg38 mapped inputs used for the human blacklist ^10^ and selected those, which showed a high cross-coverage score at a shift size equal to the read length (RSC<3.5, see Figure S11). We optimized the MACS2 q-value to maximize overlap between the EZH2_R1 and EZH2_R2 overlap, minimize the EZH2_R1 and FUS_R1 overlap, the EZH2_R1 and FUS_R2, and the percent genome covered (Table 3). The optimal MACS2 q-value was <10^-10^. We merged peaks with a distance of less than 20 kilobases, as was done for the human blacklist and removed any regions which did not show significance for at least half the inputs^10^.

### Statistical analyses

Assuming the central limit theorem^53–54^, metrics from a sufficiently large sample size (n>30) of independent identically distributed values follow the normal distribution. Otherwise, we applied the Shapiro-Wilk test ^55^ (α ≤ 0.05) to test whether calculated metrics were normally distributed. If the sample size was sufficient or the values failed to reject the Shapiro-Wilk test, parametric statistical tests were applied. To test if more than two groups’ values originate from a statistically equal population mean, an ANOVA ^56^ (α ≤ 0.001) test was applied if all group values showed a normal distribution. Otherwise, a non-parametric Kruskal-Wallis H-test ^57^ was performed and we proceeded with the analyses if the null hypothesis (no difference between the medians) was rejected. A Student t-test27^58^ was used to compare two groups from normal distributions with equal variance. A Welch t-test ^59^ was applied on to compare two groups from normal distributions with unequal variance. One sample t-tests were applied if one group value was invariant and two sample t-tests we used for all other comparisons. The Mann-Whitney U rank ^60^ was conducted on paired groups, such as metrics before and after masking artificial regions, which was not assumed to show a normal distribution. Note that to correct for the fact that the Mann-Whitney U rank compares a discrete statistic against a continuous distribution, a 0.5 continuity correction was applied to the z-score. We applied one-sided statistical tests when we expected a difference in one direction only, otherwise two-sided tests were employed. To account for multiple statistical tests p-values were adjusted using Holm’s correction^61^.

## Acknowledgements

We thank Tian Huang for help with the improved ChIP-seq pipeline development and Dr. Roberto Bonasio for comments on the manuscript. This work was supported by NSF MCB 1916431. The Arabidopsis ChIP-seq data we analyzed was from the following publication ids: PRJNA132641 ^21^, PRJNA270526 ^23^, PRJNA594407 ^20^, PRJNA560053 ^22^, PRJEB24874 ^19^, PRJEB28959 ^24^, PRJNA595112^17^, PRJNA377528 ^36^ . Cut&Run data we analyzed was from PRJNA509360^30^. The source for the twenty Arabidopsis inputs is listed in Table 1. The following twenty ENCODE input controls were used to generate the greenscreen for the hg38 genome assembly: ENCFF448TFZ, ENCFF438KJC, ENCFF880UAU, ENCFF516YKX, ENCFF349KXI, ENCFF251JQE, ENCFF495KCW, ENCFF881SJD, ENCFF433SPB, ENCFF352RKQ, ENCFF299YGP, ENCFF522TXM, ENCFF272RAI, ENCFF019HKT, ENCFF908NWF, ENCFF383ZXS, ENCFF695MWS, ENCFF048BXG, ENCFF295VUB, ENCFF476YAR.

## Competing interests

The authors declare no competing interests.

## Supplementary Tables

**Table S1.**
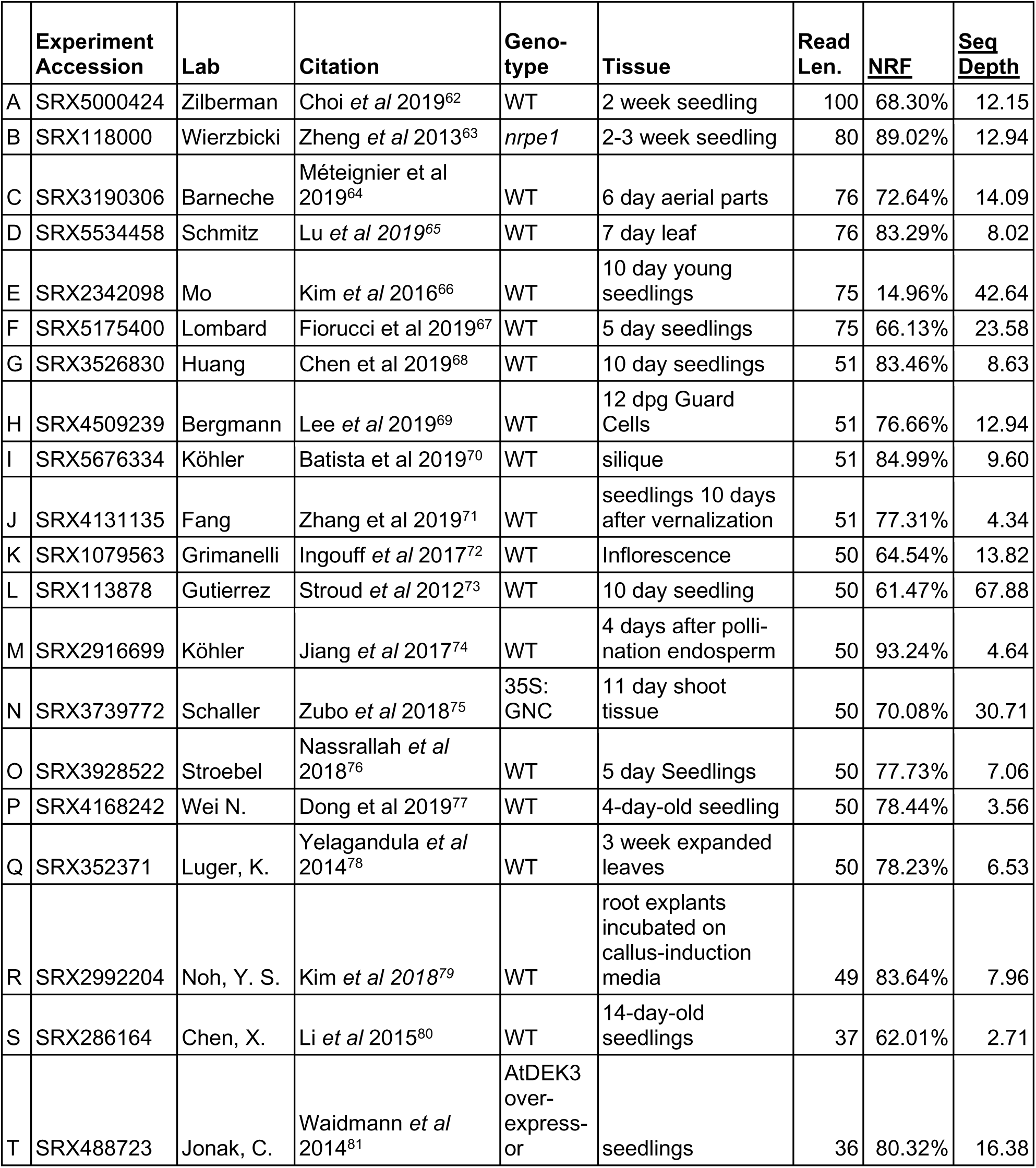
Inputs used for *Arabidopsis* Greenscreen and Blacklist. Twenty inputs generated from twenty different labs in a variety of genotypes and tissues in *Arabidopsis.* Inputs were used to identify Arabidopsis blacklist regions and greenscreen regions for ChIP-seq artifact removal (see citations for experiments in the table). Non-redundancy fraction (NRF) and sequencing (seq) depth are two metrics that measure data quality. The NRF value is the number of distinct uniquely mapping reads in a sample divided by the total number of reads. Sequencing depth, or genome coverage, is the number of reads multiplied by the read size divided by the effective genome size.

**Table S2.** Optimization of greenscreen parameters. To identify ultra-high artifacts in input experiments using greenscreen we used MACS2 to identify broad peaks, remove excess duplicate reads, and use the entire genome as background (--nolambda). We optimized the average q-value threshold (2^nd^ column) and modified the range in which regions of significance are merged (3^rd^ column). We tested these parameters by (1) quantifying enrichment of true LFY ChIP-seq peaks (overlap with LFY ChIP-chip (6^th^ column)) and (2) measuring clustering of ChIP-seq experiments for different factors (FD/TFL1 versus LFY ChIP-Seq samples; Rand index values, 7^th^ column). See Figure 3A. Finally, we minimized over-masking (percent of genome and coding genes covered; columns 4 and 5). Optimal results parameters are highlighted in green. Also shown are no mask and the Arabidopsis blacklist.

| mask type | Average<br>MACS2<br>log10<br>q-value | merge<br>distance | number of<br>basepairs<br>covered | percent<br>of<br>genome<br>covered | number of<br>genes 100%<br>covered by<br>mask | percent of LFY<br>ChIP-seq and<br>ChIPchip overlap<br>seedlings | rand index of<br>(Yk=2,<br>Y'k=2) |
| --- | --- | --- | --- | --- | --- | --- | --- |
| None | N/A | N/A | 0 | **0.00%** | 0 | 40.63% | 0.51 |
| blacklist | N/A | 5,000 | 2,854,600 | **2.39%** | 173 | 42.23% | 1.00 |
| greenscreen | 2 | 5,000 | 47,554,125 | **39.74%** | 11266 | 40.74% | 1.00 |
| greenscreen | 5 | 5,000 | 593,405 | **0.50%** | 32 | 42.40% | 1.00 |
| greenscreen | 10 | 5,000 | 484,820 | **0.41%** | 26 | 42.28% | 1.00 |
| greenscreen | 20 | 5,000 | 396,824 | **0.33%** | 25 | 42.10% | 0.85 |
| greenscreen | 30 | 5,000 | 364,173 | **0.30%** | 25 | 42.10% | 0.85 |
| greenscreen | 40 | 5,000 | 315,899 | **0.26%** | 24 | 42.00% | 0.85 |
| greenscreen | 50 | 5,000 | 311,984 | **0.26%** | 24 | 42.00% | 0.85 |
| greenscreen | 60 | 5,000 | 205,687 | **0.17%** | 17 | 41.93% | 0.85 |
| greenscreen | 10 | 0 | 129,842 | **0.11%** | 3 | 42.08% | 0.85 |
| greenscreen | 10 | 1,000 | 199,492 | **0.17%** | 6 | 42.10% | 0.85 |
| greenscreen | 10 | 2,500 | 330,118 | **0.28%** | 16 | 42.20% | 1.0 |
| greenscreen | 10 | 10,000 | 635,462 | **0.53%** | 37 | 42.28% | 1.0 |
| greenscreen | 10 | 20,000 | 958,312 | **0.80%** | 54 | 42.30% | 1.0 |

**Table S3.** Optimizing the human greenscreen mask. Basepairs and transcripts masked in the human genome by greenscreen regions (mask=GS) generated as described for Arabidopsis. Greenscreen masks derived from different input numbers (column #2) or different average maximum q-values thresholds set in MACS2 (log10(q-value), column #3), with the merge parameter set to 20kb, as for the human blacklist. Also shown are results of no mask (NM) and of the human Blacklist mask ^10^ . Parameters were optimized to cover the fewest base pairs, while retaining high peak overlap between two EZH2 replicates, low peak overlap between EZH2 (replicate 1) and FUS (replicate 1). A rand-index (Y_k=2_,Y’_k=2_) is used to quantify how well FUS, HNRNPK, and PCBP2 ChIP-seq cluster relative to biological expectation ^27^ (See Figure S5).

| mask | n inputs | avg qval limit | number of basepairs covered | percent of genome covered | Transcript mask count | total peaks in EZH2_R1 peaks | total peaks in EZH2_R1 peaks overlap EZH2_R2 | total peaks in EZH2_R1 peaks overlap FUS_R1 | Rand-index $c(Y_{k=2}, Y'_{k=2})$ |
| --- | --- | --- | --- | --- | --- | --- | --- | --- | --- |
| NM | 0 | N/A | 0 | 0.00% | 0 | 483 | 452 | 11 | .56 |
| BL | 636 | N/A | 227,162,400 | 7.36% | 3390 | 397 | 376 | 0 | 1.0 |
| GS | 20 | 2 | 57,594,369 | 1.86% | 46 | 401 | 378 | 0 | 1.0 |
| GS | 20 | 5 | 614,382 | 0.02% | 28 | 427 | 403 | 0 | 1.0 |
| GS | 20 | 10 | 354,834 | 0.01% | 20 | 431 | 407 | 0 | 1.0 |
| GS | 20 | 20 | 192,492 | 0.01% | 13 | 440 | 415 | 2 | 1.0 |
| GS | 20 | 30 | 160,668 | 0.01% | 12 | 441 | 415 | 2 | 1.0 |
| GS | 20 | 40 | 112,666 | 0.00% | 8 | 453 | 425 | 2 | .56 |
| GS | 20 | 50 | 94,070 | 0.00% | 7 | 456 | 427 | 2 | .56 |
| GS | 20 | 60 | 62,817 | 0.00% | 5 | 462 | 432 | 2 | .56 |
| GS | 10 | 10 | 340,306 | 0.01% | 27 | 431 | 407 | 0 | 1.0 |
| GS | 6 | 10 | 440,350 | 0.01% | 34 | 428 | 404 | 0 | 1.0 |
| GS | 6 | 10 | 327,769 | 0.01% | 29 | 430 | 406 | 0 | 1.0 |
| GS | 4 | 10 | 418,166 | 0.01% | 34 | 427 | 403 | 0 | 1.0 |
| GS | 3 | 10 | 332,969 | 0.01% | 26 | 433 | 408 | 0 | 1.0 |

**Table S4.**
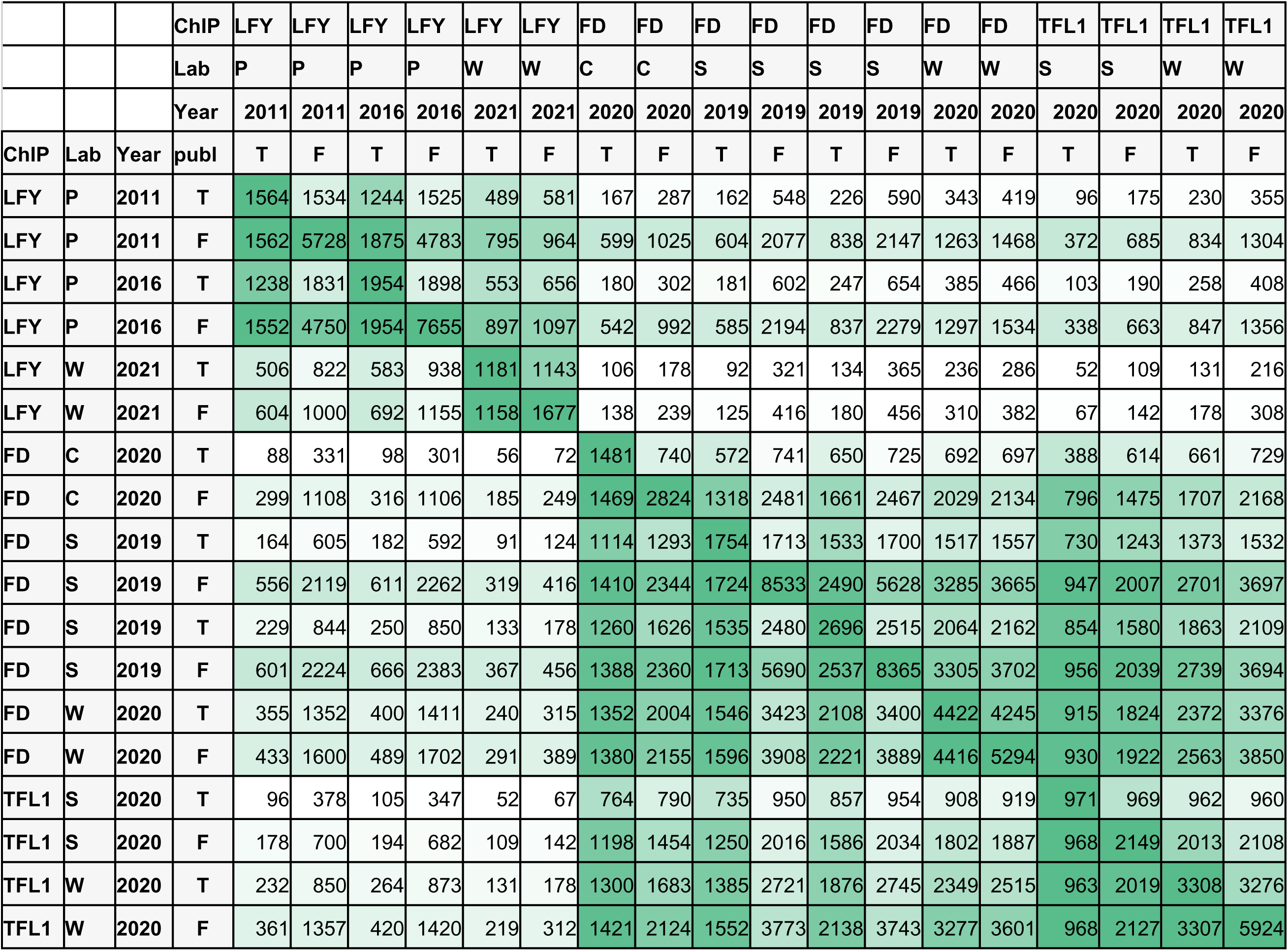
Greenscreen Peaks vs Published Peaks. Improved ChIP-seq pipelines identifies more peaks than were presented in published datasets and higher overlaps with independent ChIP-seq data for the same factor. Values in each row represent a pairwise peak overlap comparisons with ChIP-Seq samples listed in the columns. ChIP-seq samples are for two groups of factors (LFY on one hand and FD and TFL1 in the other), conducted in four labs (C=Coupland, P=Parcy, S=Schmid, W=Wagner) and published in the respective years ^17,19–24^. MACS2 controls were matched to the corresponding publications. Peaks were either identified using the pipeline discussed in this paper (publ=F) or using the methods from the respective publication (publ=T). Note that published analysis of datasets from the Wagner lab included greenscreen filters ^17,20^. See also Figure 8.

## Supplemental Figures

**Figure S1.**
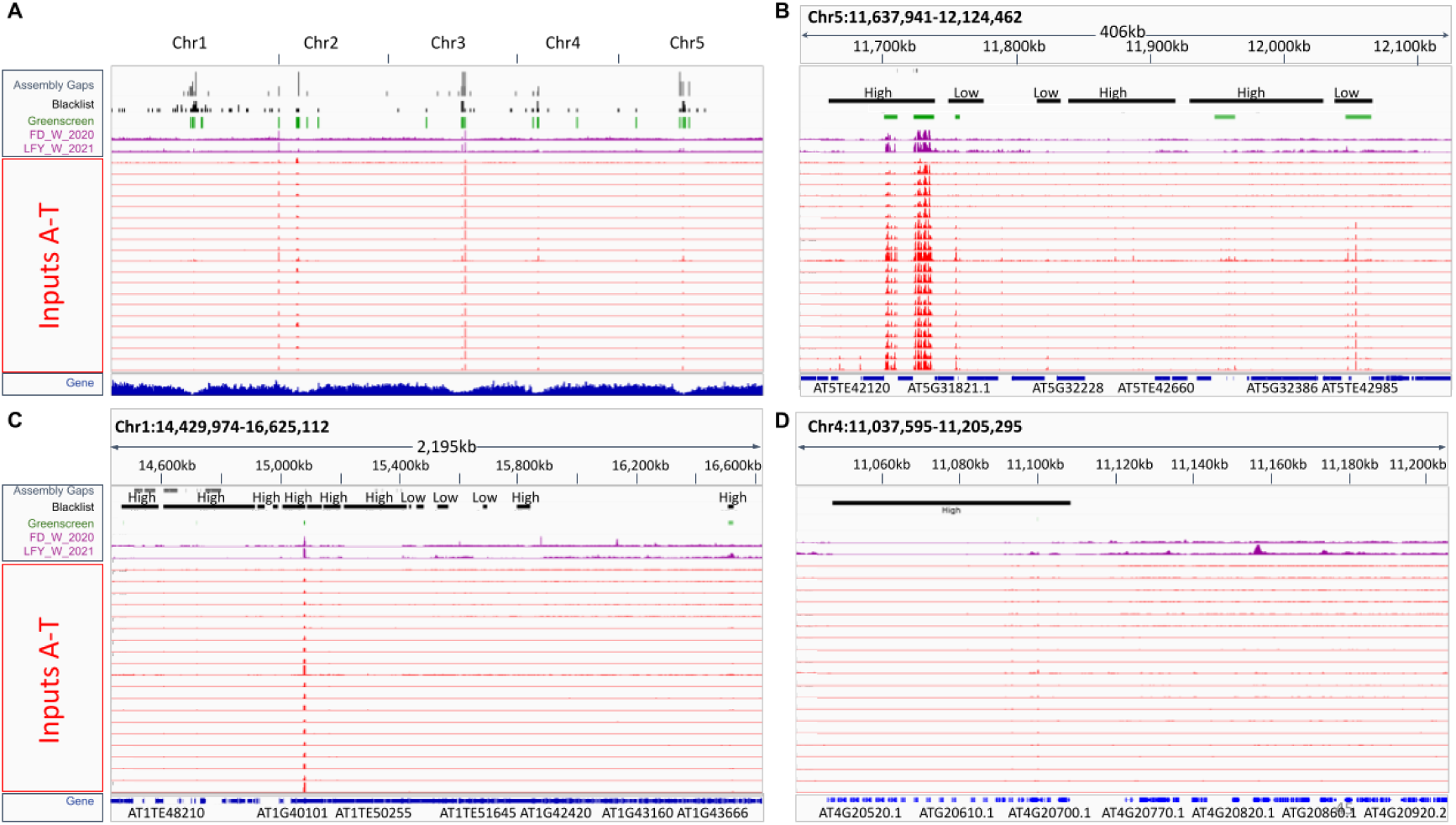
Artificial signals are present in input and ChIP samples. Integrative Genome Browser (IGV) screenshots. Top panel: indicating assembly gaps (grey), blacklist regions (black), greenscreen regions (green). LFY and FD ChIP-seq signals (purple) and twenty input control experiments (red). (A) Conserved ultra-high artifacts are readily detected in ChIP-Seq and Input-seq experiments compared to read coverage across the entire genome. (B-D) Examples of blacklist regions that have low input signals. (Scale:0-200)

**Figure S2.**
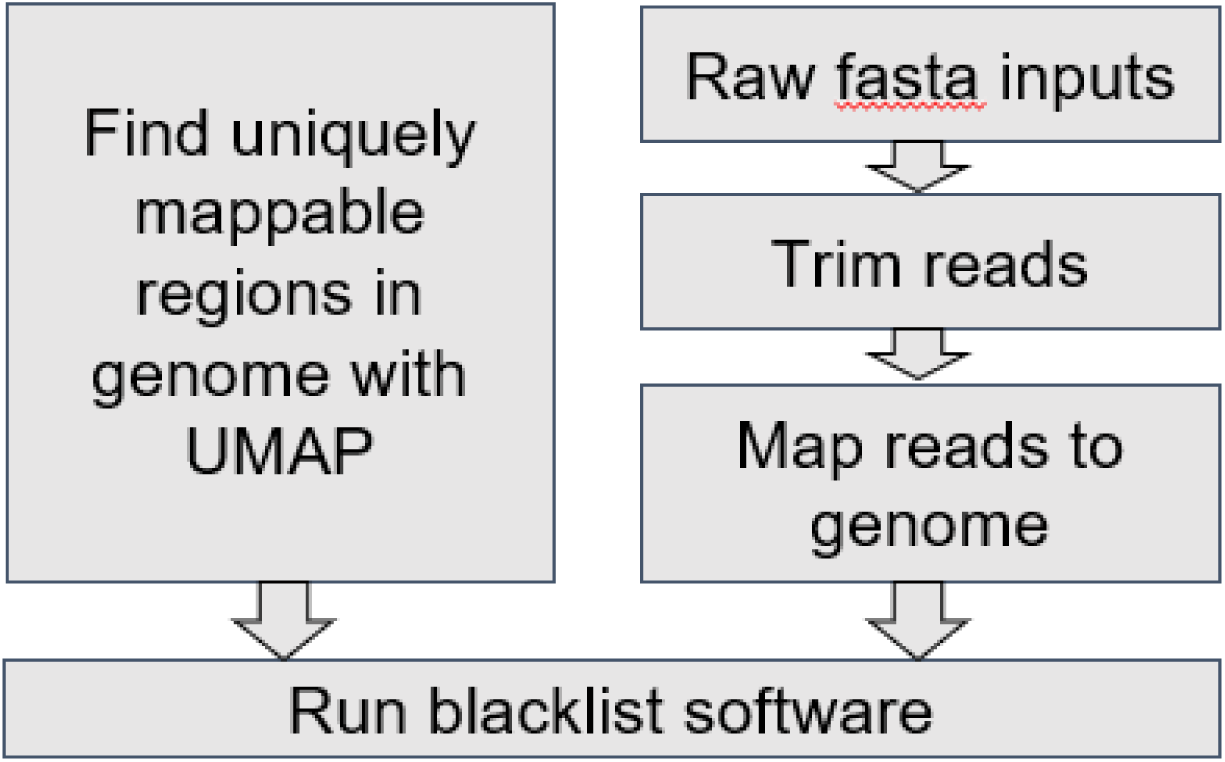
Workflow for generating a Blacklist. The blacklist software probes mappability, how well reads of specific size constraints map uniquely to genomic regions as defined by the UMAP software ^11^ and input controls to identify artifactual signal. Blacklist software uses genomic bins to quantile normalize the read depth of the inputs and then scores the bins using the median read depth value. Blacklist identifies artifact regions in the top 0.1% of signal for either input read depth or mappability, and then merges the signal regions within a set distance (20kb for human) if the top 1% of all signal is maintained or a region is unmappable.

**Figure S3.**
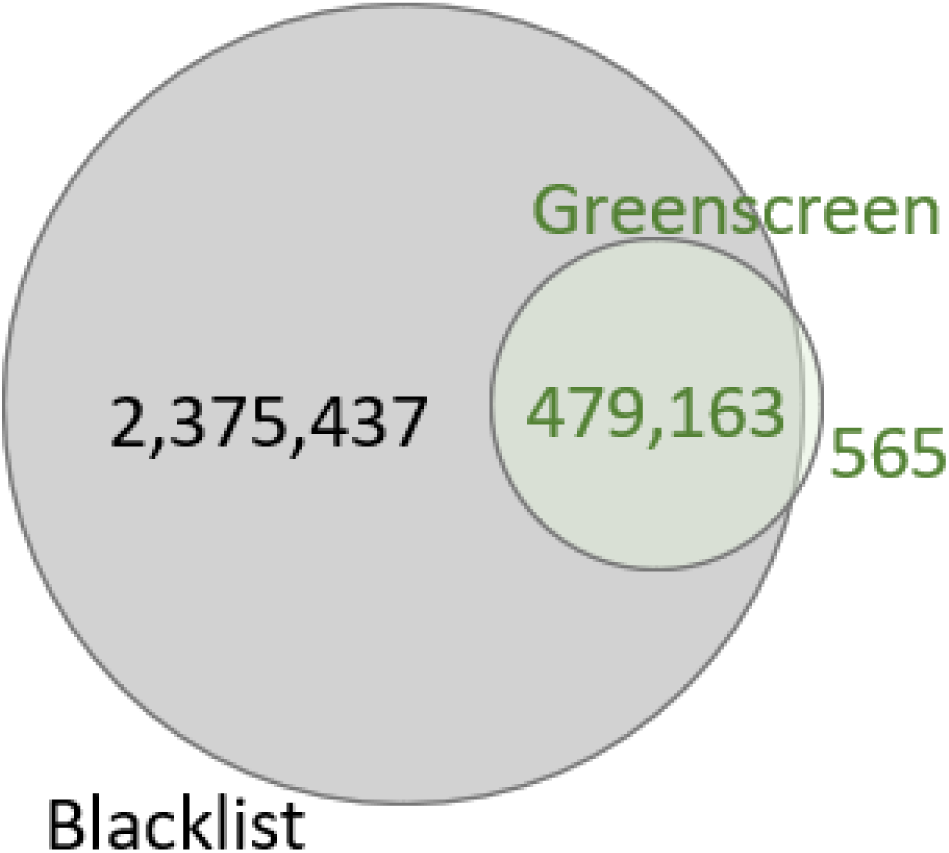
Relationship between Blacklist and Greenscreen Regions. Venn Diagram of the number of base pairs in each mask and their overlap.

**Figure S4.**
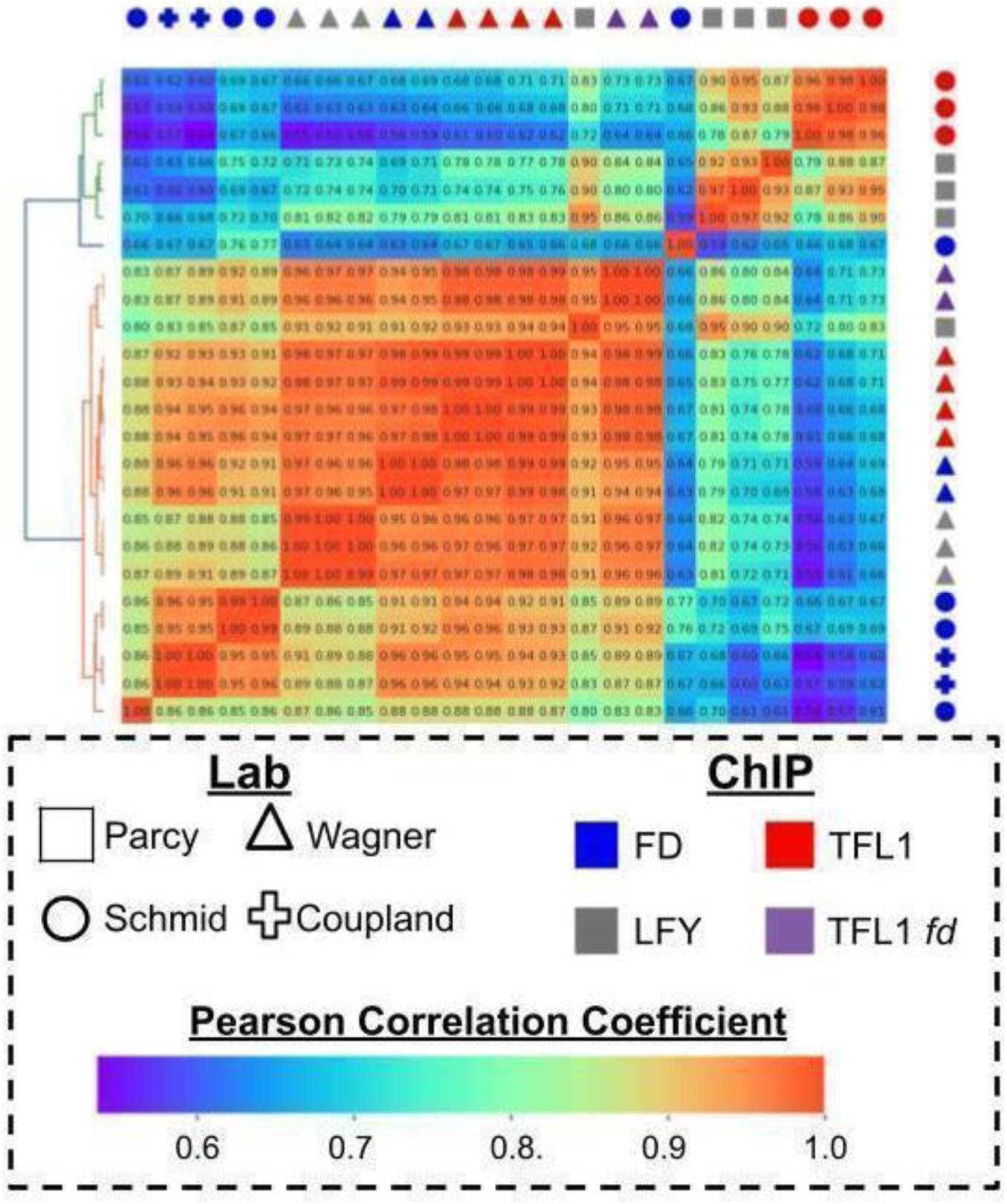
ChIP-seq artifact signal distribution. Heatmap of unsupervised clustering of artifact ChIP-seq signals in Arabidopsis greenscreen regions (left of heatmap) using pairwise Pearson Correlation Coefficients. Samples reveal spurious clustering similar to that observed in ChIP-seq samples without artifact removal.

**Figure S5.**
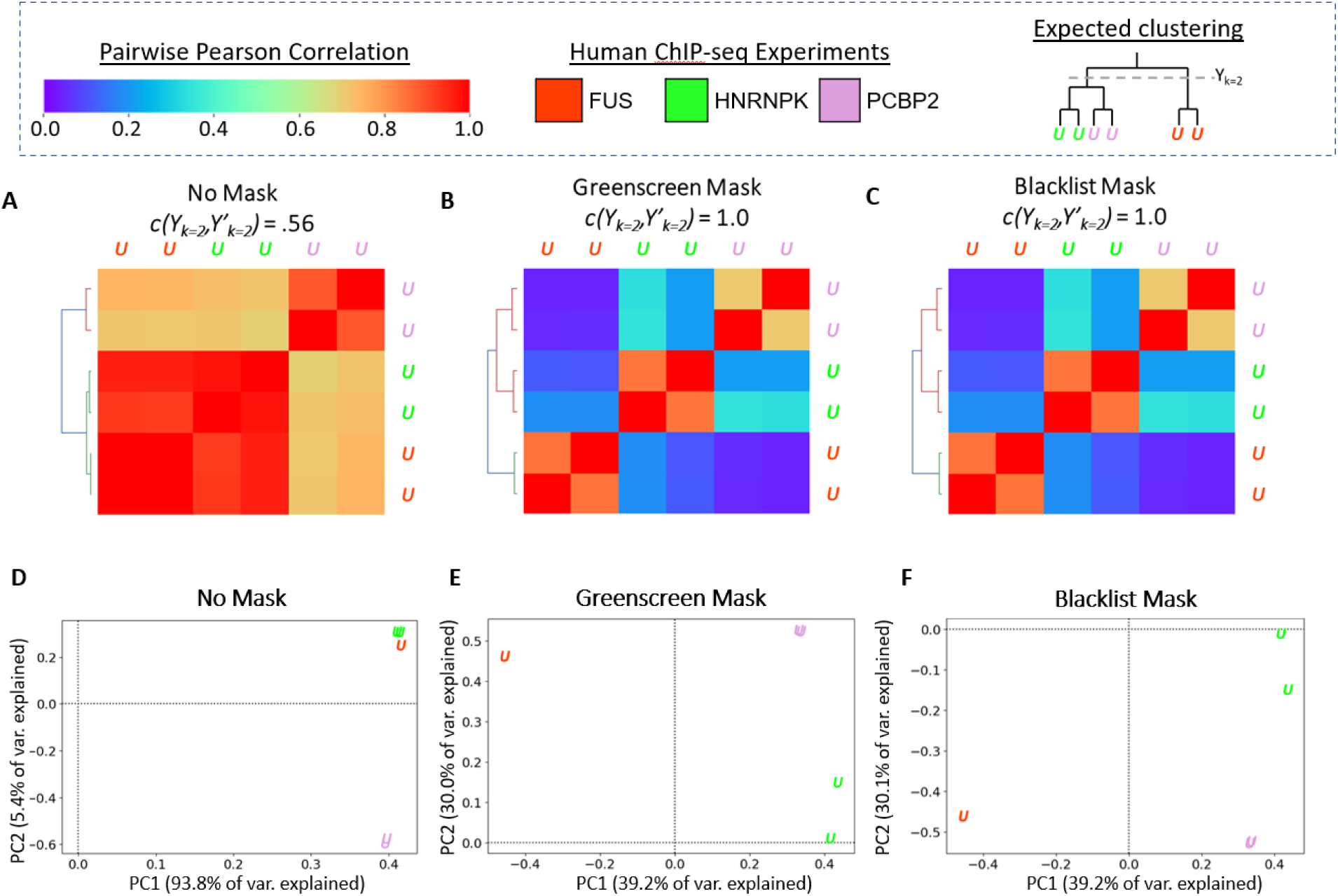
Unsupervised clustering and heatmaps of signals in FUS, HNRNKP AND PCBP2 human ChIP-seq samples. Signal in the union of peaks of FUS, HNRNPK, and PCBP2^27^ ChIP-seq replicates ^11,27^. Based on previous work it is expected that PCBP2 and HNRNPK bind more similarly than FUS ^27^. (A-C) Heatmap displays pairwise Pearson correlation coefficient values between samples signals within the peak regions. A rand-index (Y_k=2_,Y’_k=2_) is used to quantify how well the ChIP-seqs cluster compared to expectation (see right-side of legend). (D-F) Scatterplot of samples transformed using the top two principal. (A,D) no filter, (B,E) Greenscreen filter, (C,F) Blacklist filter. Color: ChIP for different factors.

**Figure S6.**
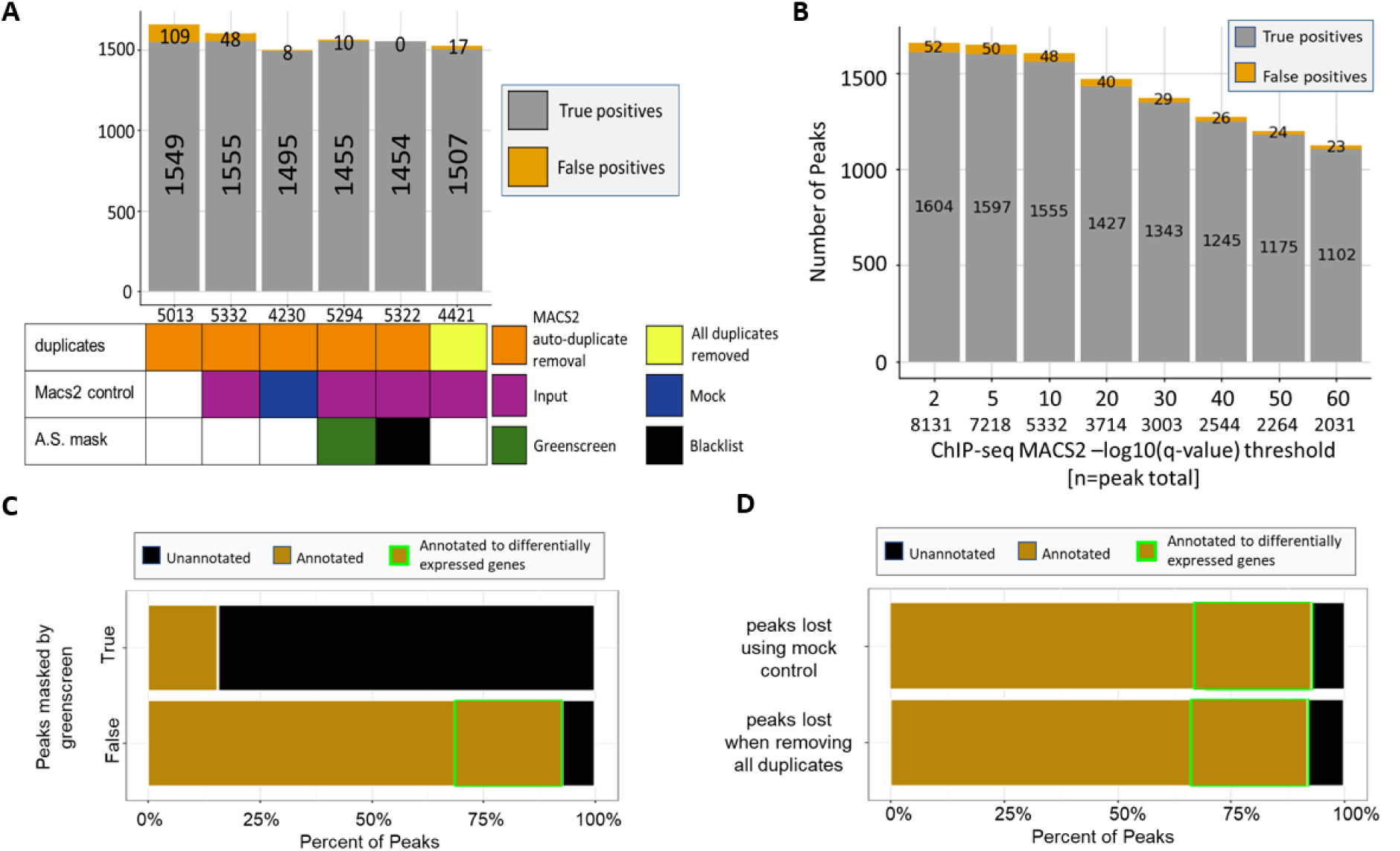
Optimizing ChIP-seq peak calling with filtering. (A) Stacked bar graphs to probe the impact of different MACS2 controls (none, mock, input), duplicate removal (MACS2 keep dup auto or no duplicates) and masking (none, greenscreen and blacklist) on calling FD ChIP-Seq MACS2 peaks ^17^. Total peak number (“n” value under x-axis), potential true positive peaks (FD ChIP-seq peaks^17^ that overlap FD ChIP-seq peaks ^19^ published by a different lab; grey bars with peak number) and potential false positive peaks (peaks overlap with the union of blacklist and greenscreen regions; orange bars with peak number). (B) Horizontal stacked bar chart to assess the impact of increasing the ChIP-seq peak summit q-value threshold for calling FD ChIP-seq peaks^17^. (C) Horizontal stacked bar charts for ChIP-seq peaks (MACS2 ‘keep dups auto’ using input controls). Top: peaks overlap with greenscreen (n=38). Bottom: peaks do not overlap greenscreen (n=5294) . Peaks were assigned to genes as described in methods; also indicated is whether genes were rapidly differentially expressed (DESeq2 adjusted p<0.005) in wild type and in *tfl1* mutants, but not *ft* mutants ^17^. (D) Horizontal stacked bar chart for FD ChIP-seq peaks ^17^ lost when using mock instead of input control in MACS2 (n=1650; top) or lost when removing all duplicates instead of applying keep dups auto in MACS2 (n=1460; bottom).

**Figure S7.**
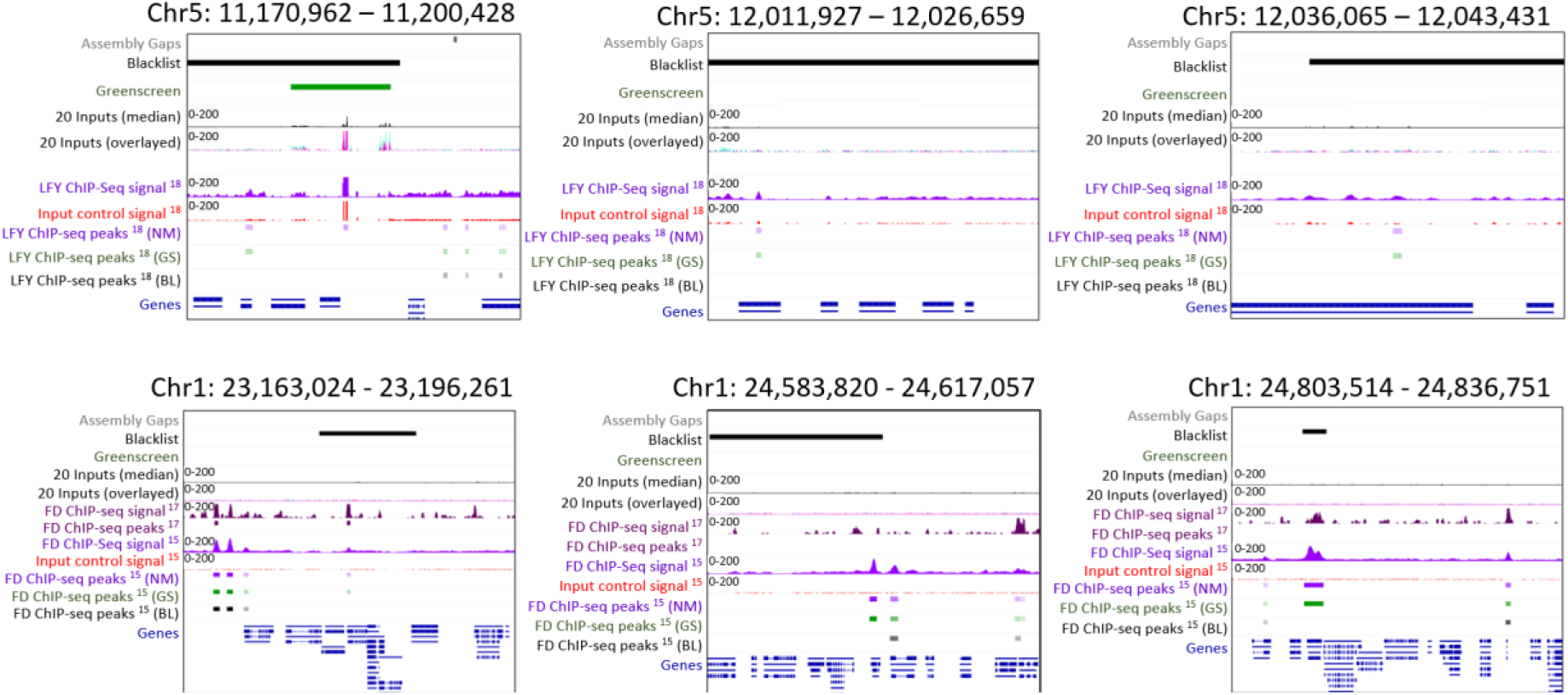
Significant peaks found in LFY and FD ChIP that overlap blacklist regions but are not masked by the greenscreen pipeline. IGV snapshots of LFY (top row) ^20^ and FD (bottom row) ^17^ peaks designated as artifacts by the blacklist but not the greenscreen filter (see also Figure 7 and Figure 9). Above: blacklist (black) and greenscreen (green) regions. Middle: median signal of the twenty inputs used to generate the blacklist and greenscreen regions and pile-up signals of the input signal in multiple colors. Bottom: independent LFY ChIP-chip peaks ^25^ and LFY ChIP-seq signals ^20^ or FD ChIP-seq signals ^17,19^ and matched input signal. Bottom and significant peaks after (NM=No Mask (purple), GS=Greenscreen Mask (green), and BL=Blacklist mask (black)). ChIP-seq and Input control signals scale: 0-200

**Figure S8.**
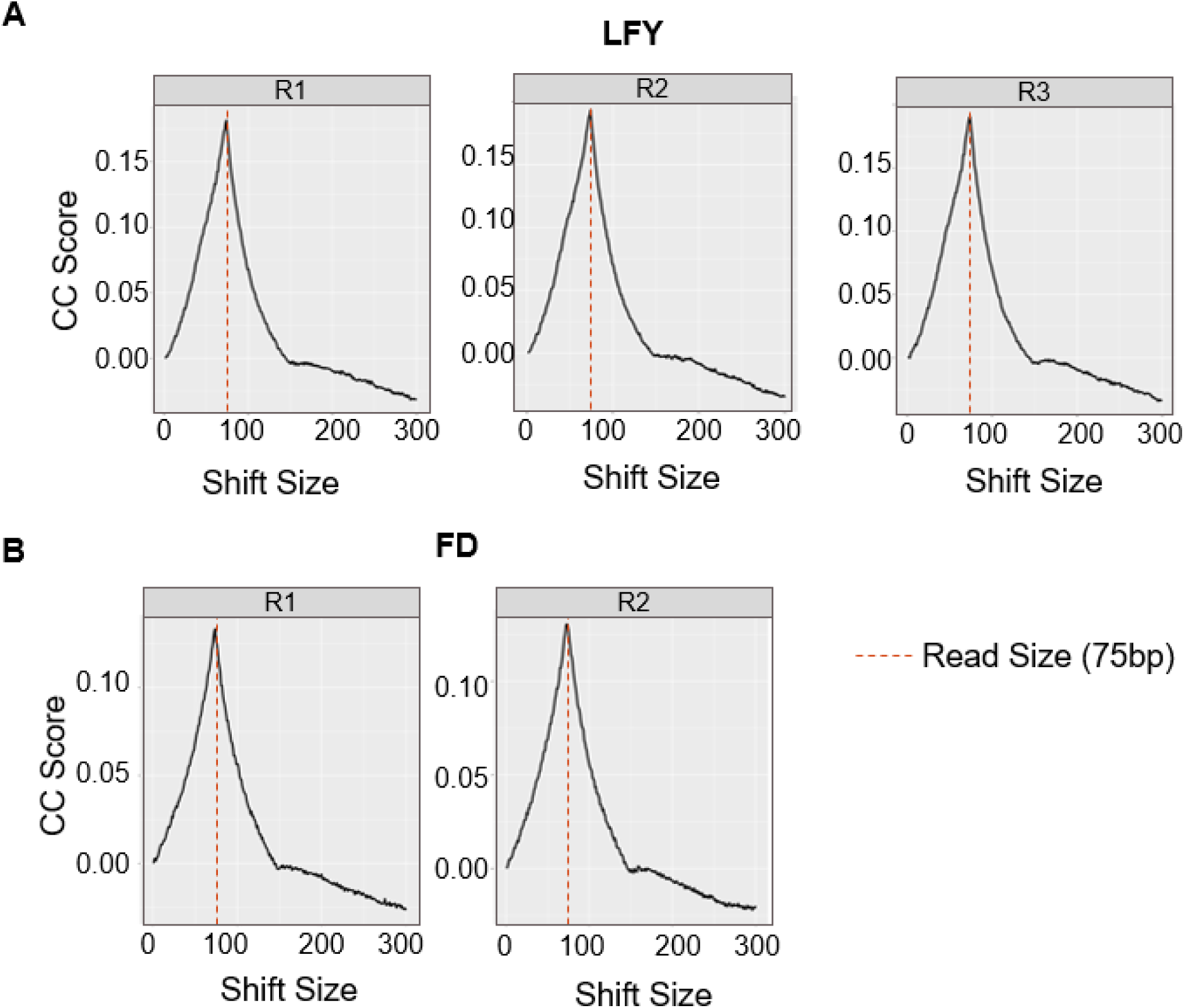
SCC plots of LFY and FD ChIP-seq reads within greenscreen regions show enrichment at a shift size equal to the read size. Cross Correlation scores of ChIP-seq reads that overlap greenscreen regions. (A) LFY ChIP-seq ^20^ and (B) FD ChIP-seq ^17^. SCC score peaks are at the read length (75bp; dashed red line).

**Figure S9.**
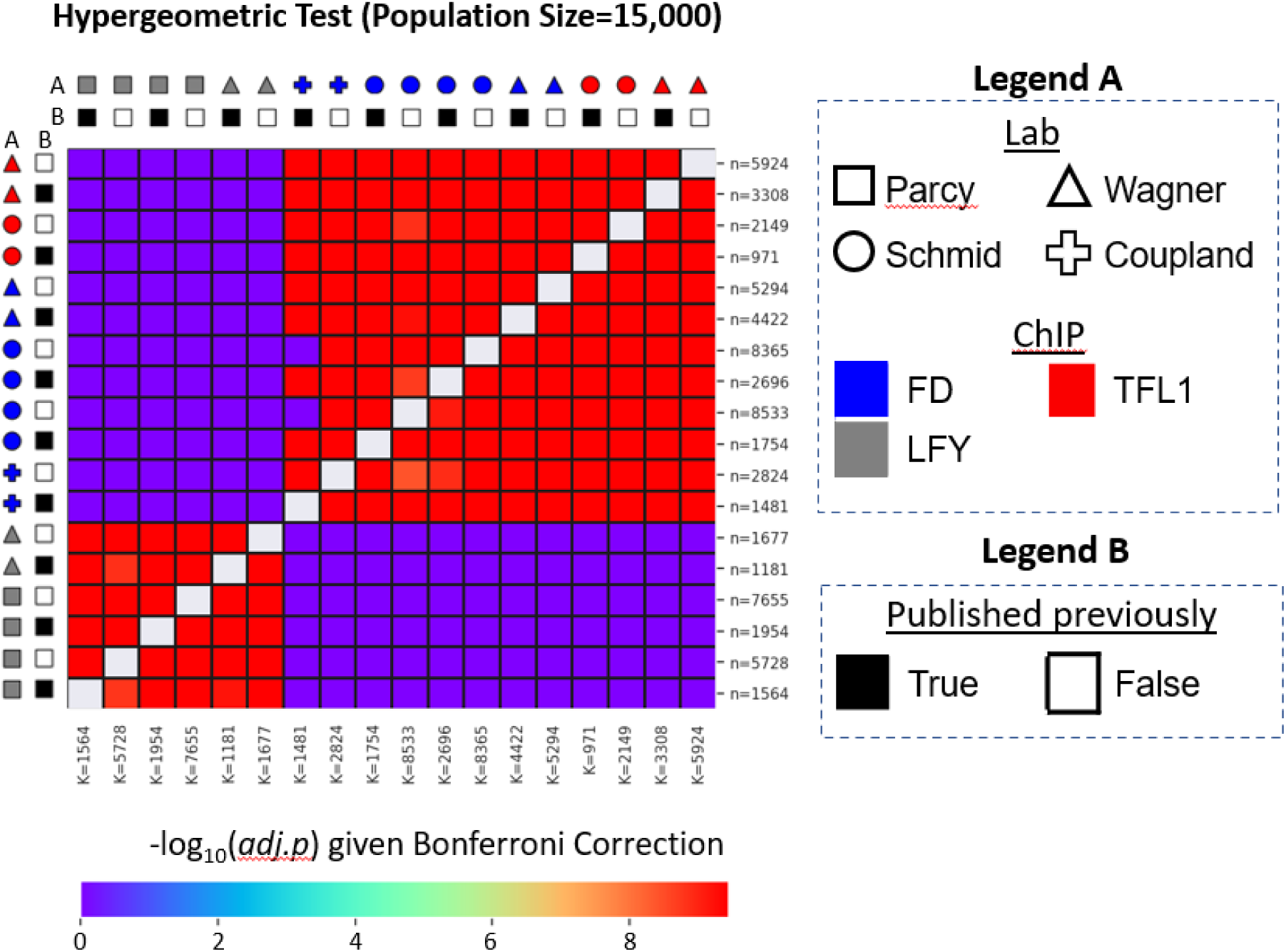
Improved ChIP-seq pipeline results in more true peaks. Hypergeometric test to measure the significance of how often peaks from one experiment (row) overlaps peaks in another experiment (column) given a population space of 15,000. Adjusted p values after Bonferroni correction. ChIP-seq samples of three factors (LFY, FD and TFL1), conducted in four different laboratories ^17,19–24^ were analyzed (Legend A). Peaks were identified using the published methods (published previously=True) or using the improved ChIP-seq method discussed in this paper (published previously=False) (Legend B). Sample size (n, listed on y-axis, number of peaks in each experiment by row), population success K (listed in x-axis, number of peaks in each experiment by column), and sample success (k, not shown, number of peaks from the experiment in a row that overlap with peaks in experiments in each column).

**Figure S10.**
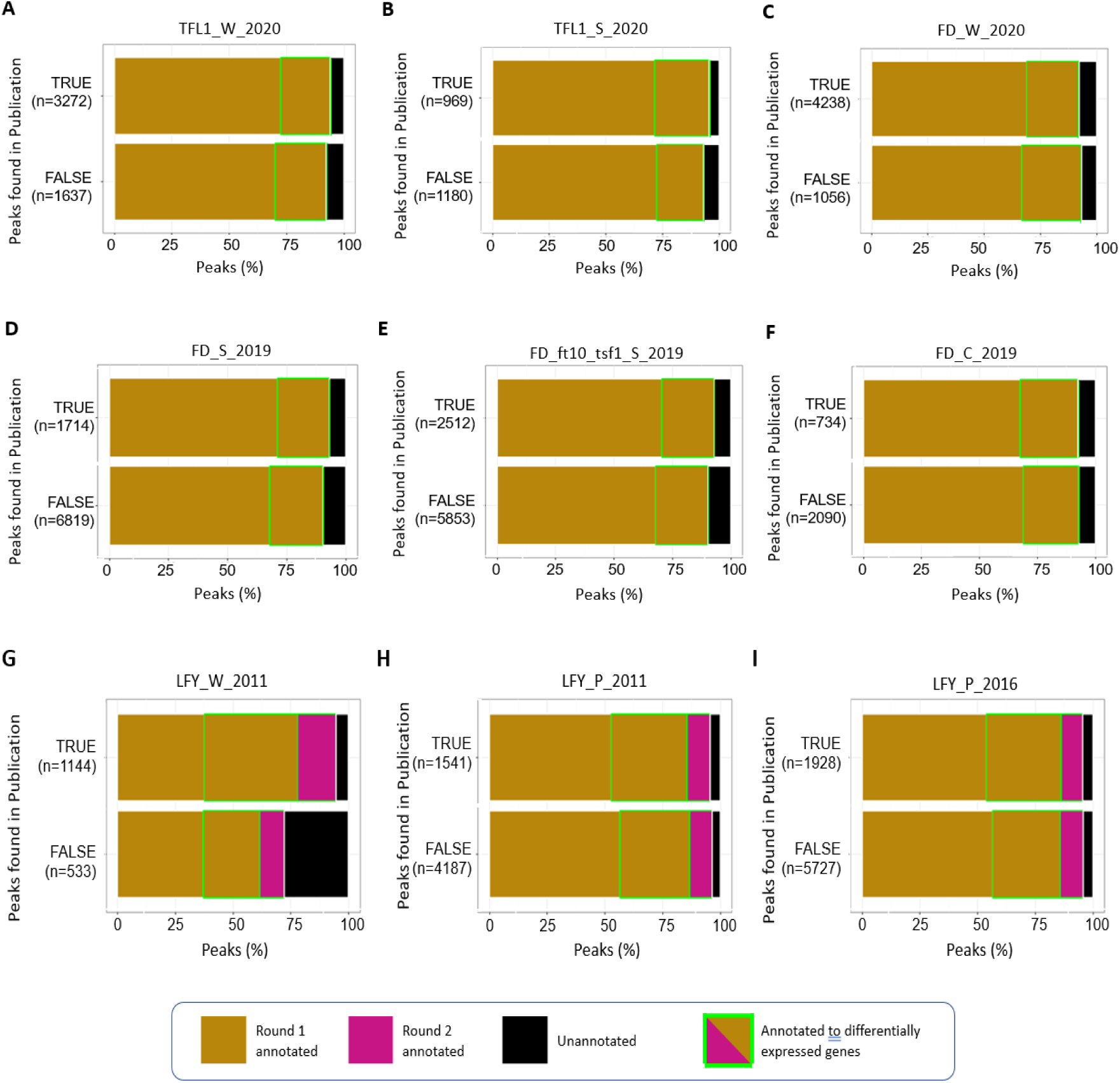
Remap 2020 hotspots that overlap with blacklist but not greenscreen regions. (A-I) Horizontal bar charts compare the new peaks found using the ChIP-seq pipeline described here to those identified in previous publications ^17,19–24^. Peaks were assigned to genes as described in methods or could not be assigned. We also indicate whether annotated genes were differentially expressed (DESeq2 adjusted p<0.005) in wild type and in *tfl1* mutants, but not *ft* mutants ^1717^ (B-G) or rapidly differentially expressed in response to either LFY factor binding ^2020^ (H-J), respectively showing in neon green box.

**Figure S11.**
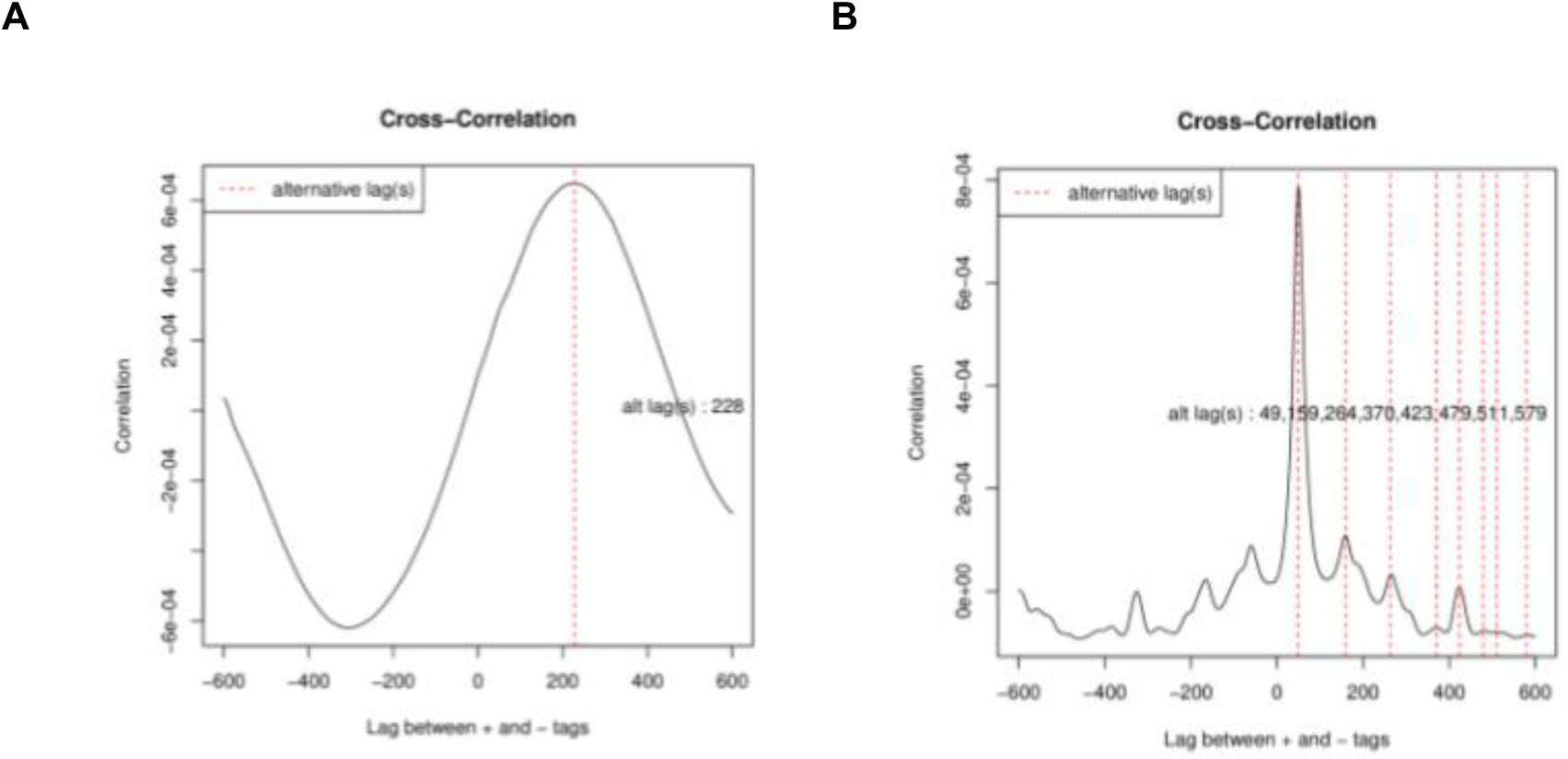
SCC profiles of published inputs. (A) An example of an input that shows enrichment at fragment size, as is expected only for ChIP-seq datasets) and was not included in the analysis ^82,35^. (B) An example of an input (Input K, Table S1) that shows enrichment only at the read length as expected for input sequences and was included in the twenty inputs used to build Arabidopsis blacklist and greenscreen filters.

## Notes

### Competing Interest Statement

The authors have declared no competing interest.

